# Clonal spreading of tumor-infiltrating T cells underlies the robust antitumor immune responses

**DOI:** 10.1101/2022.04.19.488731

**Authors:** Hiroyasu Aoki, Mikiya Tsunoda, Haru Ogiwara, Haruka Shimizu, Haruka Abe, Takaya Abe, Shigeyuki Shichino, Kouji Matsushima, Satoshi Ueha

## Abstract

The repertoire of tumor-infiltrating T cells is an emerging perspective for characterizing effective antitumor T-cell responses. Oligoclonal expansion of tumor T-cell repertoire has been evaluated; however, their association with antitumor effects is unclear. We demonstrated that the polyclonal fraction of the tumor-reactive T-cell repertoire consisting of relatively minor clones increased in tumor-bearing mice treated with anti-PD-L1 or anti-CD4 monoclonal antibody, which was correlated with antitumor effect. Meanwhile, the size of the oligoclonal fraction consisting of major clones remained unchanged. Moreover, the polyclonal fraction was enriched in progenitor exhausted T cells, which are essential for a durable antitumor response, and was more dependent on CCR7^+^ migratory dendritic cells, which are responsible for priming tumor-reactive T cells in the tumor-draining lymph nodes. These results suggest that the expansion of diverse tumor-reactive clones (“clonal spreading”) is an important mechanism by which anti-PD-L1 and anti-CD4 treatments induce robust and durable antitumor T-cell responses.

## Introduction

Immune checkpoint inhibitors (ICIs), including programmed cell death-1 (PD-1) inhibitors, are widely used to treat various types of cancer. However, only 20%–30% of patients respond to ICIs^1^, and the factors determining the differences between responders and non-responders remain unknown. The differences in clinical responses are not only due to the immune resistance mechanism of the tumor but also due to the strength of the antitumor immune response induced by ICIs^2^. Hence, characterizing the antitumor immune response that leads to better clinical outcomes is critical for optimizing patient selection and developing new therapeutic combinations.

CD8^+^ tumor-infiltrating T-lymphocytes (TILs), the primary effector cells of the antitumor immune response, are composed of various T-cell clones with different antigen specificities determined by T-cell receptors (TCRs)^3–5^. TCR transgenic mice^6^ and peptide-major histocompatibility complex (MHC) multimer technology^7^ have been used in several pre-clinical and clinical studies to analyze the characteristics of a small set of clones with high affinity to tumor antigens. More recently, TCR sequencing, a global analysis of TCRs using next-generation sequencing, has enabled the analysis of a collection of T-cell clones and the TCR repertoire^8, 9^. Hence, the TCR repertoire has been used to investigate the features of antitumor T-cell responses, leading to better therapeutic effects.

The TCR repertoire of TILs is highly skewed toward a small number of clones, in which the top 1% of the most abundant clones result in more than 20% of the TILs, suggesting the expansion of tumor-reactive clones^10, 11^. Thus, the features of the TCR repertoire of TILs have been evaluated using the clonality index^12^, representing the extent to which a part of the T-cell clones expanded in the repertoire (“clonal skewing”). Indeed, a higher clonality is associated with antitumor effects^10, 13–15^. T-cell clones that encounter antigens and proliferate in tumors differentiate into a dysfunctional state called exhaustion^16–18^. In contrast, the TIL subpopulations associated with a good response to anti-PD-1 mAb treatment are less exhausted and moderately expanded in tumors^19^. Thus, T-cell responses in which diverse tumor-reactive clones expand moderately (“clonal spreading”) are also expected to exert a desirable antitumor effect. However, the contribution of clonal spreading to the antitumor immune response has not been intensively studied.

T-cell clones that recognize epitopes unrelated to tumors also infiltrate the tumor tissue and form medium- and low-frequency clones (“bystander” clones)^20^. Thus, to assess the clonal spreading of tumor-reactive T cells, it is important to distinguish the medium- and low-frequency tumor-reactive clones from bystander clones. T-cell clones present in both tumor-draining lymph nodes (dLNs) and tumors (dLN-tumor overlapping (OL) clones) are considered to be enriched in tumor-reactive clones that are primed in the dLN and trafficked into the tumor via blood circulation^21, 22^. Considering that the newly emerged TIL clones replace exhausted clones during the anti-PD-1 therapy^23^, the overlapping TIL clones supplied by the dLNs seem to play a central role in anti-tumor T cell responses. We have previously demonstrated that an anti-CD4 monoclonal antibody (anti-CD4 mAb), which exerts potent antitumor effects in mice and humans by depleting CD4^+^ cells, including regulatory T cells (Tregs), and activating tumor-reactive CD8^+^ T-cell responses^24–27^, significantly increased the number and total frequency of dLN-tumor OL clones. However, it was unclear whether the increase in dLN-tumor repertoire overlap was also observed with PD-1 blockade and a combination of anti-CD4 and anti-PD-L1 mAbs (hereafter designated as “CD4+PDL1”), which exert synergistic anti-tumor effect^24^. Furthermore, it remains unclear whether this increase in the frequency of dLN-tumor OL clones is due to clonal skewing or spreading.

Here, we performed a T-cell receptor repertoire analysis of B16F10 and Lewis lung carcinoma tumor-bearing mice treated with anti-PD-L1 and anti-CD4 mAbs. By analyzing the dLN-tumor OL repertoire, we demonstrated that the number of TIL clones supplied by dLNs increased after anti-PD-L1 and anti-CD4 treatment owing to clonal spreading rather than clonal skewing. Moreover, the greater extent of clonal spreading in anti-CD4 and CD4+PDL1 combination treatment was correlated with their superior anti-tumor effects.

## Results

### Anti-PD-L1 and anti-CD4 treatments increased the dLN-tumor repertoire overlap in a B16 melanoma model

In our previous study, we demonstrated that anti-CD4 treatment increases the total frequency of dLN-tumor OL CD8^+^ T-cell clones^25^. Moreover, in the first-in-human trial, anti-CD4 treatment increased the tumor-associated T-cell clones in the peripheral blood, which was associated with the anti-tumor effects^27^. To examine whether the increase in the dLN-tumor repertoire overlap also occurs in anti-PD-L1 treatment or in CD4+PDL1 combination treatment, we analyzed the TCR repertoire of B16F10 tumor-bearing mice treated with anti-PD-L1 mAb, anti-CD4 mAb, and CD4+PDL1 treatments (Fig. 1a). In this model, the anti-CD4 and CD4+PDL1 treatments caused an antitumor effect on day 13, while the anti-PD-L1 treatment did not (Fig. 1b and c). The TCR repertoire of CD44^hi^ CD8^+^ T cells in the dLN and CD8^+^ T cells in the tumor was analyzed on day 14 after tumor inoculation (Fig. 1d). We then identified the dLN-tumor OL clones and compared their total frequency among the treatment groups (Fig. 1f and g). The treatment with anti-CD4 mAb and CD4+PDL1 significantly increased the total frequency of OL clones in the dLN and tumors (Fig. 1h and i). The anti-PD-L1 treatment significantly increased the total frequency of OL clones in the tumor to a lesser extent (Fig. 1i). These results suggest that both the anti-PD-L1 and anti-CD4 treatments enhanced the mobilization of tumor-reactive clones in the antitumor T-cell response.

**Fig. 1.**
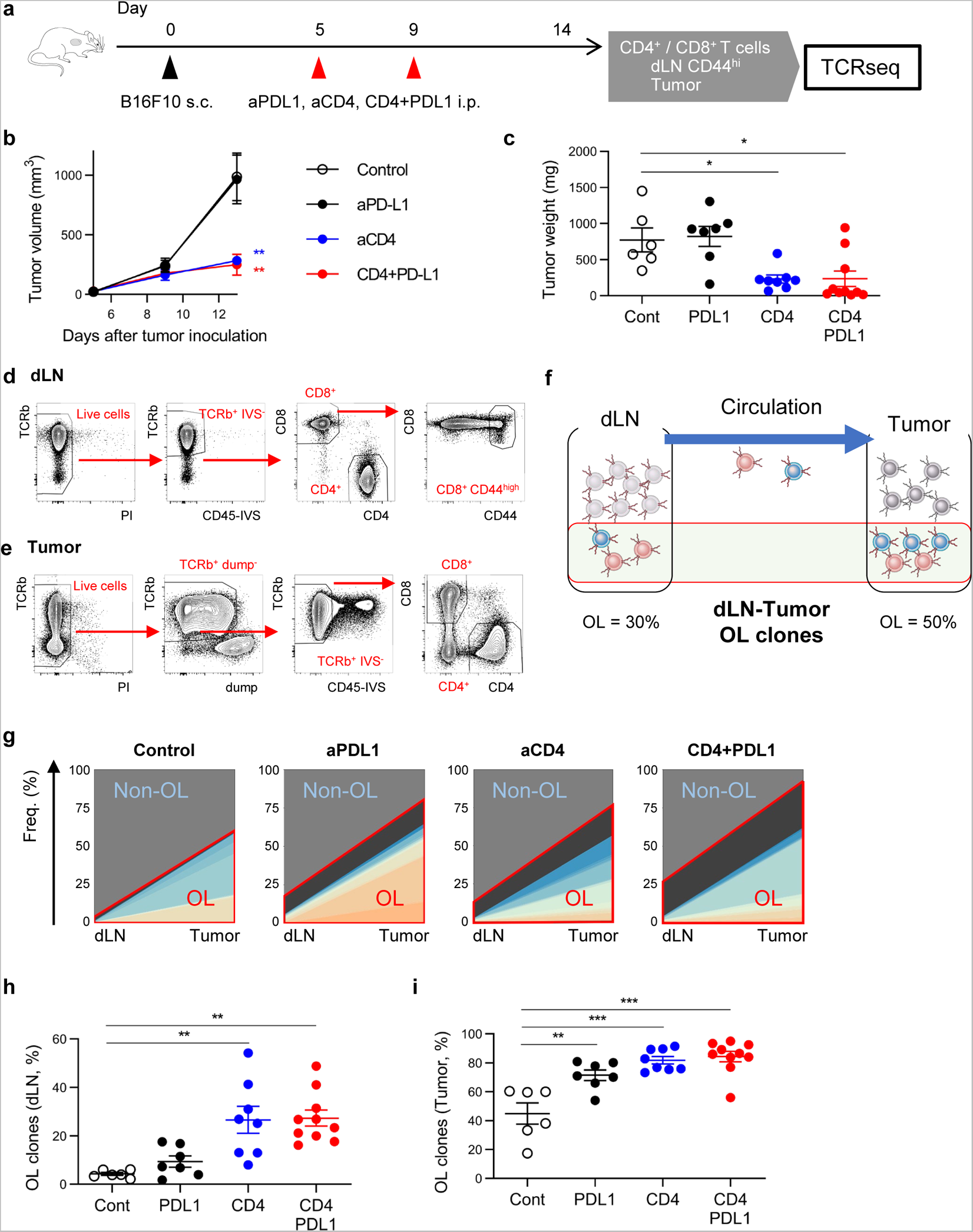
The dLN-tumor repertoire overlap increases following anti-PD-L1 and anti-CD4 treatments. **a.** Experimental procedure. **b**, **c.** Growth curve (**b**) and weight (**c**) of the tumor. **d, e.** Gating strategy for sorting the CD8^+^ CD44^high^ T cells in the dLN (**d**) and the CD8^+^ T cells in the tumor (**e**). **f.** Scheme of the dLN-tumor repertoire overlap analysis. **g.** Representative plot of the dLN-tumor repertoire overlap for each treatment group. The plot exhibits the top 20 OL clones (colored), the OL clones other than the top 20 ones (dark gray), and the non-OL clones (gray). **h, i.** Frequency of the OL clones in the dLN (**h**) and the tumor (**i**). Means ± SEM are shown (n = 6–10 per group). One-way ANOVA and Tukey’s *post-hoc* test were used in (**b**), (**c**), (**h**), and (**i**); \**P* ≤ 0.05; ** *P* ≤ 0.01; *** *P* ≤ 0.001.

The clonality of the TCR repertoire, which is the extent of clonal skewing in tissues, are commonly evaluated to monitor the antitumor T-cell responses^12^. In the dLN repertoire, clonality increased in the anti-CD4 and CD4+PDL1 groups, consistent with the increase in the dLN-tumor repertoire overlap (Supplementary Fig. 1a). In contrast, clonality did not change in the tumor repertoire between the treatment groups with different antitumor responses (Supplementary Fig. 1a). This suggests that in the B16 model, the dLN-tumor repertoire overlap reflects the effect of immunotherapies on the tumor T-cell repertoire more clearly than the commonly used clonality index.

### Clonal spreading was prominent in the dLN-tumor repertoire overlap following anti-PD-L1 and anti-CD4 treatments

Next, we examined whether the increase in the dLN-tumor repertoire overlap following the anti-PD-L1 and anti-CD4 treatments was due to clonal skewing or clonal spreading of the tumor-reactive clones. For this purpose, dLN-tumor OL clones were categorized into “top 10 clones” or “11th-or-lower clones” according to their frequency in the tumor and were considered to belong to the “oligoclonal” or “polyclonal” fraction, respectively (Fig. 2a and b). The total frequencies of oligoclonal and polyclonal overlap, which reflect the clonal skewing and spreading, respectively, were compared between the treatment groups (Fig. 2c). Oligoclonal overlap tended to increase following anti-PD-L1 or anti-CD4 treatment; however, the differences were not statistically significant. In contrast, polyclonal overlap increased significantly in the anti-PD-L1, anti-CD4, and CD4+PD-L1 groups (Fig. 2d). We also classified the clones into major (> 1%), intermediate (0.1%–1%), and minor clones (< 0.1%) based on their frequency within the tumor and confirmed that CD4 and PD-L1 expanded the polyclonal OL clones (Supplementary Fig. 1b and c). Similar results were observed in the Lewis lung carcinoma (LLC) tumor model: the polyclonal overlap increased in the treated groups, whereas the oligoclonal overlap was the same as that in the control (Supplementary Fig. 1d). These results demonstrate that the increase in the dLN-tumor repertoire overlap was mainly due to the clonal spreading of tumor-reactive T cells rather than clonal skewing.

**Fig. 2.**
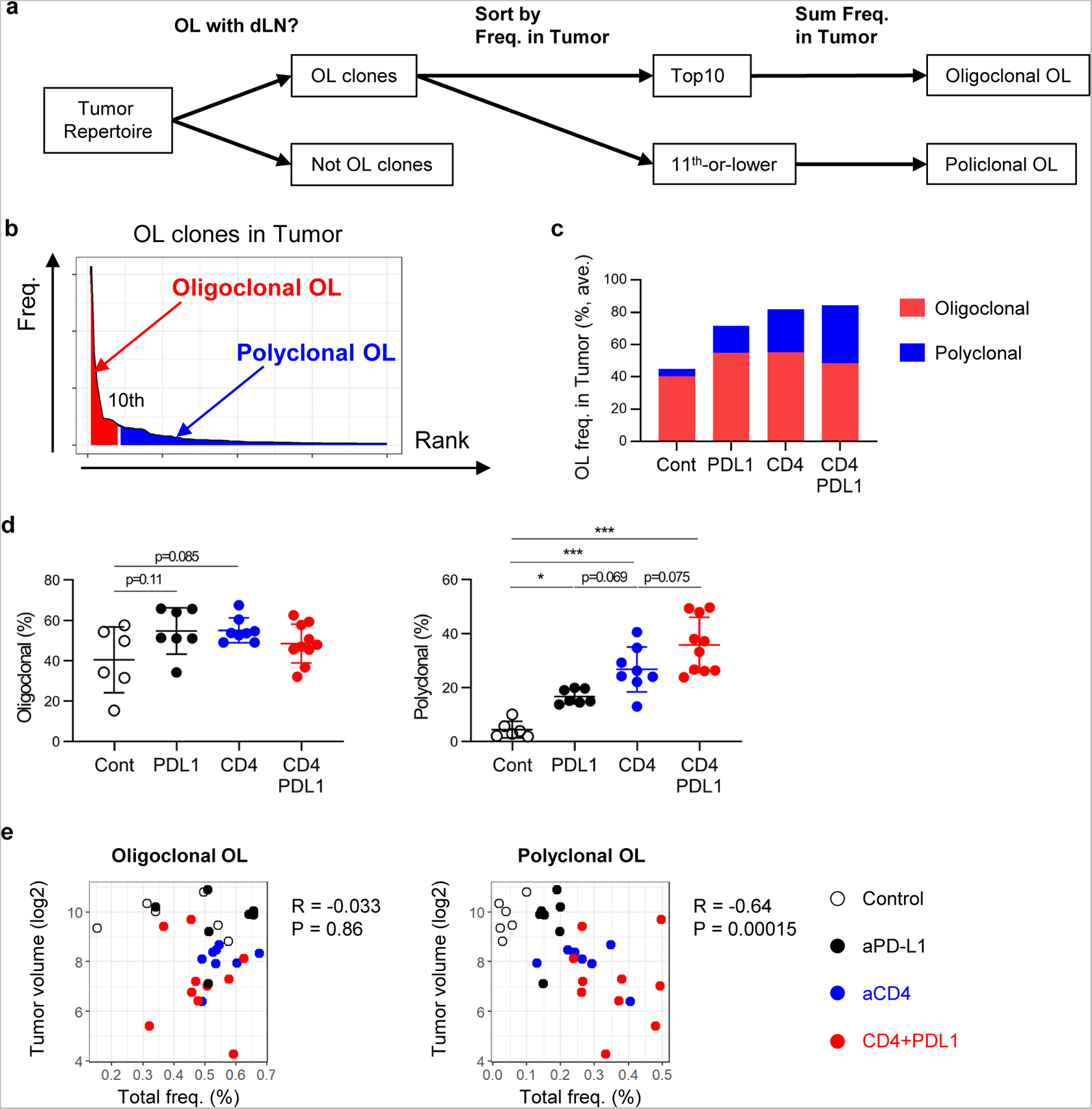
Increase in the polyclonal fraction of the dLN-tumor OL repertoire following anti-PD-L1 and anti-CD4 treatments. **a, b.** Flow chart (**a**) and representative plot (**b**) used for defining the oligoclonal and polyclonal fractions of the dLN-tumor OL repertoire. In (**b**), the frequency of individual clones is plotted according to their rank in the tumor in descending order. The top 10 (oligoclonal fraction) and 11^th^ or lower (polyclonal fraction) OL clones are colored in red and blue, respectively. **c, d.** Total frequency of the oligoclonal and polyclonal fractions of the OL clones in the tumor. **e.** Correlation between the tumor volume and the total frequency of the oligoclonal (left) or polyclonal (right) fraction of OL clones in the tumor. Means are shown (n = 6–10 per group) (**c**); means ± SEM are shown (n = 6–10 per group). One-way ANOVA and Tukey’s *post-hoc* test were used in (**d**); Spearman correlation test was used in (**e**); \**P* ≤ 0.05; *** *P* ≤ 0.001.

To clarify the association between clonal spreading and the antitumor effects of immunotherapies, we calculated the correlation between the tumor volume on day 13 and the oligoclonal or polyclonal fraction of the dLN-tumor OL repertoire (Fig. 2e). Polyclonal overlap was more strongly correlated with the tumor volume than oligoclonal overlap. A stronger correlation between the polyclonal overlap and tumor volume was also observed in the LLC model (Supplementary Fig. 1e). These results suggest that the clonal spreading of tumor-reactive T cells induced by the anti-PD-L1 and anti-CD4 treatments enhances the antitumor T-cell responses, leading to therapeutic effects.

### The polyclonal fraction of the dLN-tumor OL repertoire was enriched with progenitor-like exhausted T cells

CD8^+^ TILs display a dysfunctional state called exhaustion, which is characterized by a high expression of inhibitory receptors such as PD-1 or T-cell immunoglobulin and mucin-domain containing-3 (TIM-3)^18^. Tumors contain two subsets of exhausted T cells: terminally exhausted T cells are characterized by a high expression of inhibitory receptors and exhibit high tumor reactivity, while progenitor exhausted T cells are characterized by improved persistence and a self-renewal capacity^28–31^. Because progenitor exhausted T cells can proliferate and generate terminally exhausted ones following PD-1 blockade, this subset is thought to be essential for a durable antitumor T-cell response^29^. Based on these findings, we then investigated whether the progenitor exhausted cells corresponded to a polyclonal fraction of the dLN-tumor OL repertoire, which reflects clonal spreading.

To answer this question, single-cell targeted RNA sequencing and TCR sequencing (scRNA/TCRseq) were performed using a BD Rhapsody system (Fig. 3a). CD8^+^ TILs were collected from B16F10 tumor-bearing mice treated with anti-CD4 mAb, and a library was generated for scRNA/TCRseq. Out of the 12,397 cells analyzed, paired TCR sequences were detected in 10,649 cells (pairing efficiency, 85.9%). Eight clusters were identified based on the single-cell gene expression profiles (Fig. 3b and Supplementary Fig. 2a). Based on the expression levels of marker genes (Fig. 3c and Supplementary Fig. 2b) and the gene expression signature scores (Fig. 3D and Supplementary Fig. 2c), we characterized these clusters as “naïve (NV)-like” (#7), “progenitor exhausted” (#4 and 6), and “terminally exhausted” (#0, 1, 2, 3, and 5). Among the “progenitor exhausted” clusters, cluster 4 showed an intermediate level of exhaustion scores and a lower expression level of *Tcf7*, suggesting an intermediate state between progenitor exhausted and terminally exhausted states (Fig. 3c and d). These findings were supported by the result of pseudotime analysis, which revealed the trajectory from cluster 7 (NV-like) through clusters 4 and 6 (progenitor exhausted) to clusters 0 and 1 (terminally exhausted) (Supplementary Fig. 2b). Among the “terminally exhausted” clusters, cluster 2 was highly proliferative and cluster 3 was more cytotoxic, according to the *Gzma* expression level and the cytotoxicity score (Supplementary Fig. 2c and d). Moreover, cluster 5 was characterized by a high expression of interferon-responsive genes, such as *Isg15* and *Mx1* (Fig. 3c and Supplementary Fig. 2d). The clonality of the “terminally exhausted” clusters was higher than the “progenitor exhausted” and “NV-like” clusters, showing that the “terminally exhausted” clusters were composed of highly expanded clones (Supplementary Fig. 2e).

**Fig. 3.**
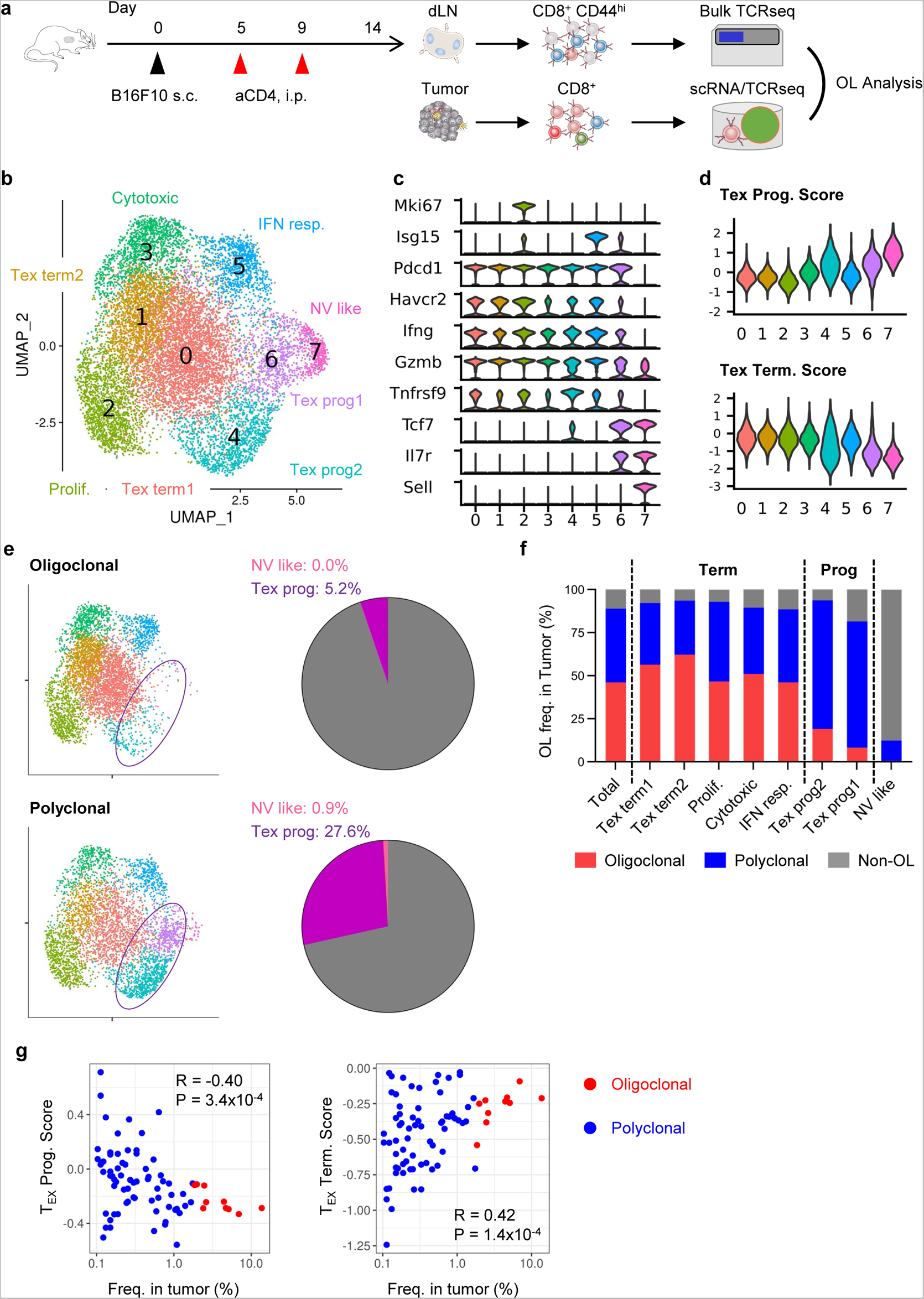
Single-cell analysis of the dLN-tumor OL clones following anti-CD4 mAb treatment. **a.** Experimental procedure. **b.** Uniform manifold approximation and projection (UMAP) plot of the scRNA-seq data of CD8^+^ TILs following anti-CD4 treatment. **c.** Violin plot showing the expression of marker genes in the T-cell subsets. **d.** Violin plot showing the module scores of the feature genes of progenitor (top) and terminally (bottom) exhausted T cells. **e.** UMAP showing the T cells in the oligoclonal or polyclonal fraction of the dLN-tumor OL repertoire (left). Pie chart showing the proportion of T cells in the “naïve-like” (pink) or “progenitor exhausted” (purple) clusters in the oligoclonal or polyclonal fraction (right). **f.** Bar chart showing the proportion of the oligoclonal (red) and polyclonal (blue) fractions of the OL clones and the non-OL clones (gray) in each cluster. **g.** Correlation between the clone’s frequency in the tumor and the progenitor or terminally exhausted score of each clone. The progenitor or terminally exhausted score was calculated for each cell as in (**d**), and the average value was then calculated for each clone. Pearson correlation test was used in (**e**); Tex: terminally exhausted; Tex int: intermediately exhausted; Tex prog: progenitor exhausted; IFN resp.: IFN responding; NV-like: naïve-like; Prolif.: proliferating.

To compare the phenotype of the T-cell clones consisting of the oligoclonal and polyclonal fractions of the dLN-tumor OL repertoire, we identified the OL clones using the results of bulk TCR-seq performed on dLN CD8^+^ CD44^high^ T cells collected from the same mouse (Fig. 3a). Oligoclonal or polyclonal OL clones were projected onto a uniform manifold approximation and projection (UMAP) plot (Fig. 3e, left). The oligoclonal and polyclonal fractions of the dLN-tumor OL repertoire contained very few cells from the “NV-like” cluster (#7), which supported the tumor reactivity of the OL clones. The proportion of “terminally exhausted” clusters (#0, 1, 2, 3, and 5) was over 50% in the oligoclonal and polyclonal overlap. At the same time, the proportion of “progenitor exhausted” clusters (#4 and 6) was higher in the polyclonal fraction (5.2% in the oligoclonal fraction vs. 27.6% in the polyclonal fraction; Fig. 3e, right). Consistently, the polyclonal fraction was predominant in the “progenitor exhausted” clusters (Fig. 3f and Supplementary Table S1). Moreover, the genes representing the progenitor- and terminally exhausted phenotypes were highly expressed in the polyclonal and oligoclonal fraction, respectively (Supplementary Fig. 2f). At the clone level, a higher frequency of OL clones in the tumor was associated with a lower score for the progenitor exhausted state and a higher score for the terminally exhausted state (Fig. 3g). These results demonstrate that the oligoclonal OL mainly consisted of terminally exhausted T cells, while the polyclonal fraction of the dLN-tumor OL repertoire was enriched in progenitor exhausted T cells.

### The polyclonal fraction of the OL repertoire was dominant in the Ly108^+^ Tim-3^−^ TIL subpopulation

To confirm the results of the scRNA/TCRseq analysis in a high-throughput manner, we performed bulk TCR repertoire analysis on the CD8^+^ TIL subpopulations at different exhaustion stages. Using Tim-3 and Ly108 as cell surface markers for progenitor and terminally exhausted T cells^29, 31^, the CD8^+^ TILs from B16F10 tumor-bearing mice treated with anti-PD-L1 and anti-CD4 were sorted into PD1^−^, PD1^+^ Ly108^+^ Tim-3^−^ (“Ly108”), PD1^+^ Ly108^+^ Tim-3^+^ (“DP”), and PD1^+^ Ly108^−^ Tim-3^+^ (“Tim-3”) subsets using flow cytometry (Fig. 4a). Among these subpopulations, T-cell exhaustion is thought to progress in the following order: PD1^−^, Ly108, DP, and Tim-3^29^. We then analyzed the TCR repertoire of these four TIL subsets as well as CD44^hi^ CD8^+^ T cells in the dLN. The clonality of the TCR repertoire was the lowest in PD1^−^ and gradually increased in the DP and Tim-3 subsets in anti-CD4-treated mice (Fig. 4b). A similar tendency was observed in the anti-PDL1 and CD4+PDL1 groups (Supplementary Fig. 3a). These results show that the repertoire of the PD1^−^ and Ly108 subsets was more polyclonal than that of DP and Tim-3, consistent with the scRNA/TCRseq analysis results (Supplementary Fig. 2d). Next, we attempted to verify whether the polyclonal overlap predominated in the progenitor exhausted population. For this purpose, we estimated the frequency of each clone in the total CD8^+^ TILs (“pooled”) by estimating the frequency within the TIL subsets (PD1^−^, Ly108, DP, and Tim-3) weighted by the total cell count of these subsets (Fig. 4c, example). We confirmed that the frequency of each clone in the pooled repertoire, which was estimated as described previously, was correlated with the frequency in the real CD8^+^ TIL repertoire (Supplementary Fig. 3b). Subsequently, the OL clones in the oligoclonal and polyclonal fractions were identified based on their frequency in the pooled CD8^+^ TIL repertoire (Fig. 4c), and their total frequency in each T-cell subset was calculated (PD1^−^, Ly108, DP, and Tim-3). In the anti-CD4 group, the proportion of the oligoclonal fraction was significantly higher in the DP and Tim-3 subsets than in the PD1^−^ and Ly108 subsets, while that of the polyclonal fraction was the highest in the Ly108 subset (Fig. 4d). Consistently, the proportion of the polyclonal fraction surpassed that of the oligoclonal fraction in Ly108 cells (Fig. 4e). These results confirmed the findings of the scRNA/TCRseq analysis that the oligoclonal fraction predominated among terminally exhausted T cells, while the polyclonal fraction contained more progenitor exhausted T cells in the dLN-tumor OL repertoire. Notably, the CD4+PDL1 group exhibited results similar to those of the anti-CD4 group; however, this tendency was not observed in the anti-PDL1 group (Supplementary Fig. 3c and d). This may reflect a weaker degree of clonal spreading in the anti-PDL1 group than in the anti-CD4 or CD4+PDL1 group.

**Fig. 4.**
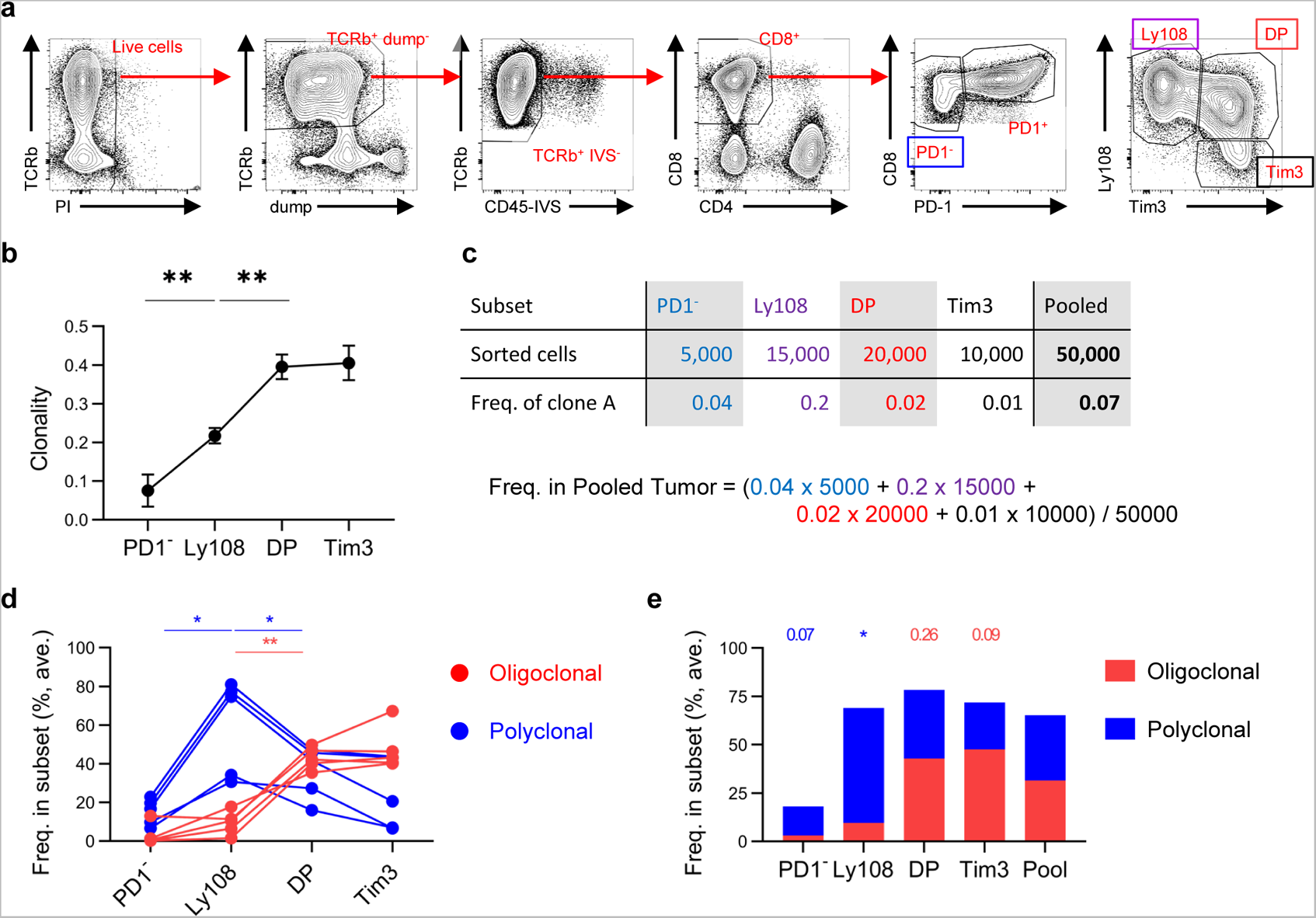
Oligoclonal and polyclonal fractions of the dLN-tumor OL repertoire in CD8^+^ TIL subsets following anti-CD4 mAb treatment. **a.** Gating strategy used for sorting the PD1^−^, Ly108, DP, and Tim-3 CD8^+^ TIL subsets in anti-CD4 treated mice. **b.** Clonality of the CD8^+^ TIL subsets. **c.** Example of the estimation of clone frequency in pooled CD8^+^ TILs based on the frequencies in each CD8^+^ TIL subset. **d.** Frequency of the oligoclonal (red) or polyclonal (blue) fraction of OL clones in the CD8^+^ TIL subsets. Lines indicate the data from the same mice. **e.** Bar chart showing the proportion of the oligoclonal (red) or polyclonal (blue) fraction of the OL clones in each CD8^+^ TIL subset. Mean ± SEM is shown in (**b**) and the mean is shown in (**e**) (n = 5 per group) (**b**); one-way RM ANOVA and Tukey’s *post-hoc* test were used in (**b**) and (**d**); paired t-test was used in (E); \**P* ≤ 0.05; ** *P* ≤ 0.01.

### Enhanced priming by migratory DCs in the dLN contributed to clonal spreading following anti-PD-L1 and anti-CD4 treatments

Our results from the TCR repertoire analysis proposed that the clonal spreading of tumor-reactive T cells contributed to the antitumor effect induced by anti-PD-L1 and anti-CD4 treatments in the B16F10 model. As for the mechanism of clonal spreading, we hypothesized that the anti-PD-L1 and anti-CD4 treatments enhanced the priming of more diverse T-cell clones in the dLN. Migratory DCs in the dLN, which deliver tumor antigens and play a critical role in inducing tumor-reactive CD8 T-cell responses^32, 33^, express a higher level of PD-L1 than resident DCs. Moreover, because Tregs deprive co-stimulatory molecules on migratory DCs through CTLA-4^34^, the anti-CD4 treatment, which removes CTLA-4 expressing Tregs, seems to enhance the priming of CD8^+^ T cells by migratory DCs in the dLN. To examine whether migratory DC-dependent priming contributes to the clonal spreading after the treatment, we utilized Fscn1^DTR-Cre^ knock-in mice to remove the CCR7^+^ FSCN1^+^ migratory DCs in the dLN (Supplementary Fig. 4a). FSCN1 is an actin-bundling protein that promotes the migration of DCs into LNs^35–37^. Since FSCN1 is expressed not only in CCR7^+^ migratory DCs but also in neurons^35, 38^, we inoculated B16F10 cells into FSCN1 KI bone marrow chimeric mice (FSCN1 BMC). In tumor-bearing FSCN1 BMC mice, DTR was weakly expressed on a part of CCR7^+^ MHC II^+^ DCs in the tumor and strongly expressed on most migratory DCs in the dLN (Supplementary Fig. 4b). Administration of DT to these mice slightly decreased the number of CCR7^+^ MHC II^+^ DCs in the tumor and completely depleted migratory DCs in the dLN (Supplementary Fig. 4c).

We subjected B16F10-bearing FSCN1 BMC mice to anti-PD-L1 and anti-CD4 treatments, with or without DT administration (Fig. 5a). Depletion of migratory DCs in the dLN of DT-treated mice was confirmed on day 14 (Fig. 5b-d). TCR repertoire analysis revealed that the total frequency of dLN-tumor OL clones in the tumor decreased in mice receiving DT in the anti-PD-L1 and CD4+PD-L1 groups (Fig. 5e). The oligoclonal fraction of the dLN-tumor OL repertoire was equivalent or higher in DT-treated mice compared to DT-untreated mice (Fig. 5f). In contrast, the polyclonal fraction of the dLN-tumor OL repertoire decreased following DT administration in all three treatment groups (Fig. 5g). These results suggest that the anti-CD4 and anti-PD-L1 antibodies enhance the priming of tumor-reactive T-cell clones by migratory DCs in the dLN, leading to clonal spreading in tumor-bearing mice.

**Fig. 5.**
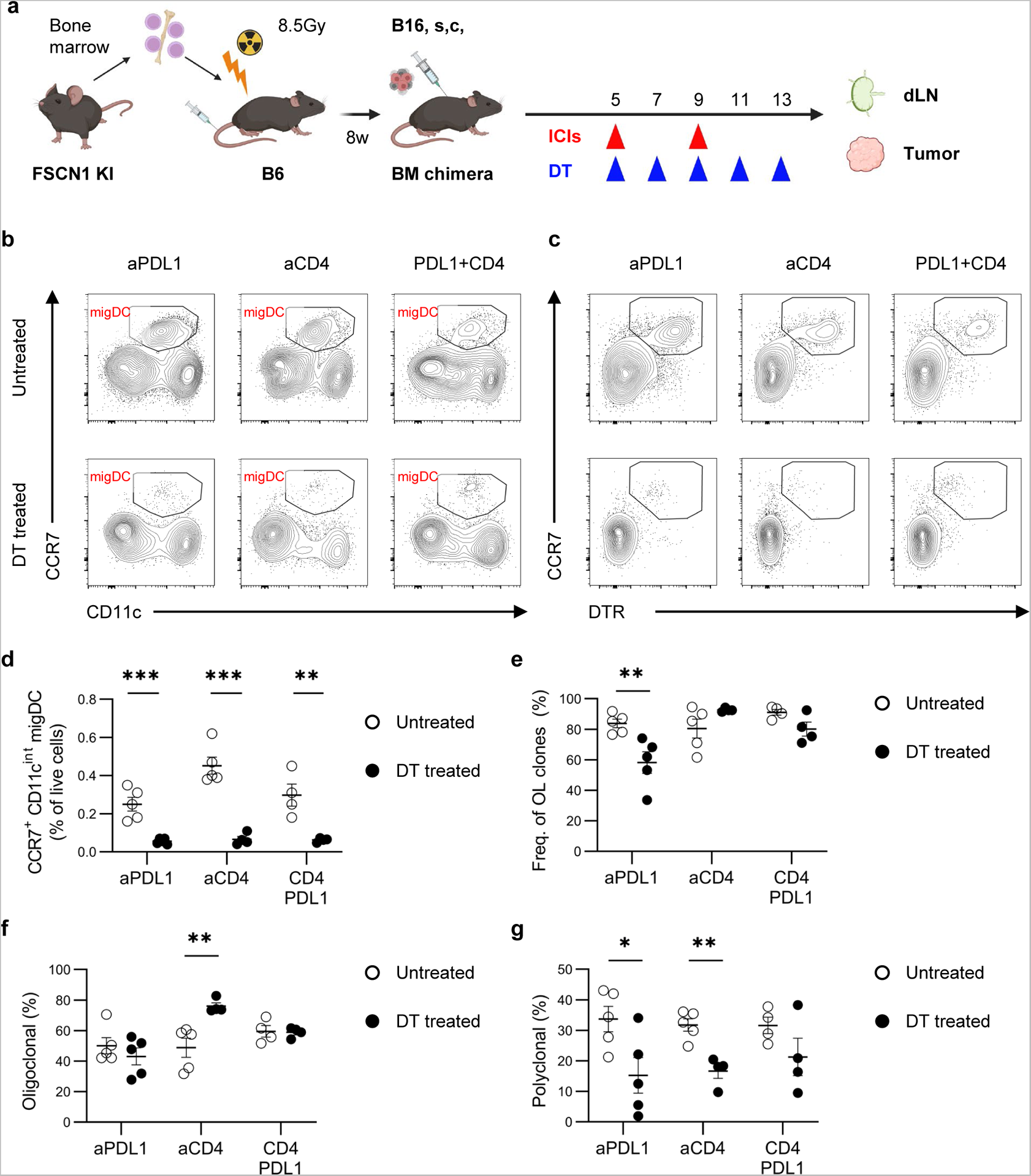
Decrease in the polyclonal fraction of the dLN-tumor OL repertoire by depleting migratory DCs following anti-PD-L1 and anti-CD4 treatments. **a.** Experimental procedure. **b, c.** Representative flow cytometry plot of the dLN lineage^−^cells for CCR7^+^ CD11c^int^ (**b**) and CCR7^+^ DTR^+^ (**c**) migratory DCs (migDCs) in untreated or DT-treated mice on day 14. **d.** Comparison of the frequency of CCR7^+^ CD11c^int^ migDCs in dLN between DT-treated and untreated mice. **e.** Comparison of the frequency of OL clones in the tumor between DT-treated and untreated mice. **f, g.** Comparison of the frequency of the oligoclonal (F) and polyclonal (G) fraction of dLN-tumor OL clones in the tumor between DT-treated and untreated mice. **a.** was created using BioRender.com; means ± SEM are shown (n = 4–5 per group); multiple unpaired t-tests were used in (**d**), (**e**), (**f**) and (**g**); \**P* ≤ 0.05; ** *P* ≤ 0.01; *** *P* ≤ 0.001.

## Discussion

We demonstrated that the anti-PDL1 and anti-CD4 mAb treatment increased the CD8^+^ T-cell repertoire overlap in the dLN and tumor, mainly owing to clonal spreading. In the B16F10 model, anti-CD4 and CD4+PD-L1 treatments exert strong antitumor effects whereas anti-PD-L1 treatment alone shows a limited antitumor effect^24^. In the present study, the antitumor effect of anti-CD4, anti-PD-L1, and CD4+PD-L1 treatments correlated with the extent of clonal spreading. Single-cell analysis of the gene expression and TCR sequences of TILs revealed that progenitor exhausted T cells, which are essential for a durable antitumor response to PD-1 blockade^29–31^, were enriched in a polyclonal fraction of the OL repertoire, which reflected clonal spreading. Furthermore, enhanced priming in the dLN by migratory DCs contributed to the clonal spreading of tumor-reactive clones following anti-PD-L1 and anti-CD4 treatments. These results suggest that clonal spreading induces antitumor T-cell responses by anti-PD-L1 and anti-CD4 treatments in the B16 melanoma model.

Our finding that clonal spreading were enhanced after anti-PD-L1 and anti-CD4 treatments seems to contradict with many previous studies on the repertoire analysis of TILs reporting an association between oligoclonal expansion and objective antitumor effects following ICIs^10, 13–15^. However, these studies did not separate CD4^+^ and CD8^+^ TILs prior to TCR sequencing. CD8^+^ TILs exhibit higher clonality than CD4^+^ TILs^19^ in most cases, and the clonality of the total TIL repertoire is reported to be correlated with the proportion of CD8^+^ cells^39^. This evidence suggests that the higher clonality in these studies reflects a larger proportion of CD8^+^ cells in TILs. Here, we performed a TCR repertoire analysis of purified CD8^+^ T cells and clarified the clonal composition changes in the CD8^+^ T-cell repertoire under immunotherapy, which was not evident in previous studies.

TCR repertoire analysis on dLN-tumor repertoire overlap clearly demonstrated that the anti-PD-L1 and anti-CD4 treatments did not increase clonal skewing but enhanced the clonal spreading of tumor-reactive T cells. Notably, the increase in the polyclonal fraction of the TIL repertoire following anti-PD-L1 and anti-CD4 treatments was not observed when non-overlapping clones were included in the analysis. Thus, the analysis of the dLN-tumor OL, which enriches the TIL clones supplied by dLNs, is important for quantifying the clonal spreading of tumor-reactive T cells and evaluating antitumor T cell responses.

scTCR analysis of TILs and bulk TCR repertoire analysis of exhausted TIL subsets revealed the immunological phenotypes of the oligoclonal and polyclonal fractions of the dLN-tumor OL repertoire; unfortunately, we could not analyze untreated mice owing to the low density and number of CD8^+^ TILs in the B16F10 model. Recent studies on the scTCR analysis of human TILs reported that highly expanded CD8^+^ T cell clones exhibited a terminally exhausted phenotype with tumor reactivity^16, 40, 41^. Consistent with these findings, the oligoclonal fraction of the dLN-tumor OL also largely consisted of T cells with a terminally exhausted phenotype and contained a small amount of progenitor exhausted T cells. Conversely, the proportion of progenitor exhausted T cells in the polyclonal fraction of dLN-tumor OL was higher than that in the oligoclonal fraction. In addition, Naïve-like and PD1^−^ CD8^+^ TILs with low tumor-reactivity^5^ were enriched in the non-OL repertoire but not in the polyclonal fraction of OL, as expected. Considering that terminally exhausted T-cell clones inside the tumor were replaced following PD-1 blockade in cancer patients^23^, the OL clones in the polyclonal fraction may function as reserves while oligoclonal fraction may not persist for a long period. Furthermore, it is also presumed that clonal spreading owing to anti-PD-L1 and anti-CD4 treatments enlarges the reservoir of tumor-reactive T-cell clones, which underlies the robust and persistent antitumor effect.

One question that remained unaddressed was how the anti-PD-L1 and anti-CD4 treatments enhance the clonal spread of tumor-reactive T cells. We initially hypothesized that the enhanced priming of tumor-reactive clones in the dLN increases the polyclonal fraction of the dLN-tumor OL repertoire. Indeed, in mouse models, the activation of T cells in the dLN is enhanced by anti-CD4^24^ and anti-PD-L1 treatments^42^. The priming of tumor-reactive T cells in the dLN is mainly mediated by CCR7^+^ migratory DCs, which capture tumor antigens, mature in the tumor, and migrate to the dLN via lymphatics^32, 33^. Migratory DCs express a higher level of PD-L1, whose interactions with PD-1 on activated CD8^+^ T cells is inhibited by PD-1 blockades in the dLN^42^. Moreover, the anti-CD4 treatment, which removes CTLA-4 expressing Tregs, may promote the priming of CD8^+^ T cells through increasing the expression of CD80/86 on migratory DCs in the dLN. These observations support the possibility that clonal spreading is promoted by the enhanced priming capacity of migratory DCs. In line with this, the specific depletion of migratory DCs in FSCN1^DTR-Cre^ KI BMC mice reduced the polyclonal fraction of the dLN-tumor OL repertoire. Our results underscore the contribution of the enhanced priming in the dLN to the clonal spreading of tumor-reactive clones.

This study had some limitations. First, we could not demonstrate the relationship between the extent of clonal spreading and clinical outcomes in patients with cancer under ICI treatment. This is attributable to our limited access to the dLN specimen. TCR repertoire analysis on a bilateral subcutaneous tumor model, in which tumor cells are inoculated bilaterally into the backs of mice, could be an alternative approach that provide the evidence for a causal relationship between clonal spreading and the antitumor effect of ICIs^43, 44^. Second, we could not analyze the immunological differences between clones contributing to clonal skewing and clonal spreading, including the tumor reactivity, TCR affinity, and exhaustion status. Clones in oligoclonal OL, which are observed in tumor-bearing mice, regardless of the treatment, are thought to exhibit higher affinity for cancer antigens and are able to expand under immune checkpoints but are more prone to exhaustion. Conversely, clones in polyclonal overlap, whose expansion is enhanced by ICIs, are thought to have TCRs with lower affinity for cancer antigens and need more costimulatory signals for expansion^45^. Cloning the TCR alpha- and beta-chain pairs using scTCR sequencing and generating TCR transgenic T cells of various dLN-tumor OL clones would enable us to validate the features and functions of T-cell clones contributing to clonal skewing or clonal spreading. Here, we defined the oligoclonal/polyclonal fraction based on the rank or frequency of clones in the tumor. However, a more refined definition, which incorporates the immunological features of clones, such as the exhaustion status or TCR affinity, can more precisely evaluate the extent of clonal spreading and become a more reliable marker for the treatment outcome.

In summary, our results suggest that an enhanced clonal spreading underlies the robust antitumor immune responses by which anti-PD-L1 and anti-CD4 treatments. Overall, we emphasize the importance of evaluating the polyclonal nature of antitumor T-cell responses through TCR repertoire analysis, which provides a novel index that reflects the efficacy of ICI treatments.

## Methods

### Materials availability

The Fscn1 knockin mouse (Accession No. CDB0153E: https://large.riken.jp/distribution/mutant-list.html) will be distributed upon requests. All antibodies, reagents, oligonucleotides, and software and algorithms used in this study are listed in Supplementary Table S2-5, respectively.

### Data and code availability

All sequencing data (Bulk TCRseq, and Single-cell RNA/TCR sequencing) have been deposited at GEO and are publicly available as of the date of publication (GEO: GSE198211).

All original code has been deposited at Github (https://github.com/hiro-aoki-mriid/Aoki_ICIrepertoire).

### Mice

Eight-week-old female C57BL/6 mice (CD90.2) were purchased from Sankyo Labo service corporation inc. CD90.1 congenic mice with gp100 melanoma antigen-specific Pmel-1 TCR transgene (Pmel Tg, #:005023) were purchased from The Jackson Laboratory, then B6 mice were backcrossed for at least 6 generations. Female Pmel Tg mice were used for the experiment at 8-10 weeks old. All mice were bred at specific pathogen-free facilities in Research Institute for Biomedical Sciences, the Tokyo University of Science and in RIKEN Kobe branch.

### Cell lines

Lewis lung carcinoma (LLC) was kindly provided by Dr. Fuminori Abe (Nippon Kayaku). B16F10 is a gp100^+^ spontaneous murine melanoma cell line, kindly provided by Dr. N. Restifo (National Cancer Institute, MD). Both cell lines were cultured in RPMI-1640 medium (Nakarai) supplemented with 10% fetal bovine serum (MP Bio Japan), 1% HEPES (Nakarai) and 1% penicillin/streptomycin (Nakarai) at 37°C in a humidified atmosphere at 5% CO_2_.

### Generation of Fscn1DTR-Cre knock-in mice

The Fscn1 knockin mouse (Accession No. CDB0153E: https://large.riken.jp/distribution/mutant-list.html) was generated by CRISPR/Cas9-mediated knockin in zygotes as previously described ^46^. For the homologous recombination-mediated knockin, the donor vector consisting of homology arms and P2A-DTR-P2A-Cre (insert gene) was generated to insert the P2A-DTR-P2A-Cre cassette at the 3-base upstream of the PAM sequence. For microinjection, the mixture of crRNA (CRISPR RNA) (50ng/ul), tracrRNA (trans-activating crRNA) (100ng/ul), donor vector (10ng/ul) and TrueCut™ Cas9 Protein v2 (100ng/ul, ThermoFisher) were injected into the pronucleus of one-cell stage zygote from C57BL/6 mice. From 233 zygotes, 57 F0 mice were obtained and 3 of them were knockin mice confirmed by PCR, in which the PCR fragments of KI allele with 5’FW and 5’REV primers (1301 bp), and 3’FW and 3’REV primers (587 bp) were detected (Supplementary Fig. 4a). The germline transmissions of knockin F0 mice were confirmed by genotyping of F1 mice.

### Generation of BM chimeric mice

Recipient B6 mice were lethally irradiated (8.5 Gy, split into 2 doses given 3 hours apart) the day before transplantation with 3 × 10^6^ Fscn1-KI BM cells. Mice were used for the experiments at least 8 weeks after transplantation.

### In vivo treatment

B16F10 or LLC cells (5 × 10^5^ cells except for Supplementary Fig. 3b: 2 × 10^5^ cells) were inoculated subcutaneously (s.c.) into the right flanks of C57BL/6 or Fscn1-KI BMC mice. Anti-PD-L1 mAb (clone 10F.9G2, BioLegend) or Anti-CD4 mAb (clone GK1.5, BioLegend) was injected intraperitoneally (i.p.) at a dose of 200 μg per mouse on days 5 and 9 after tumor inoculation. DT (FUJIFILM Wako) was injected intraperitoneally (i.p.) at a dose of 25 ng/g continuous every second day until day 5. Tumor diameter was measured twice weekly and used to calculate tumor volume (mm^3^) [(major axis; mm) x (minor axis; mm)^2^]. All animal experiments were conducted in accordance with institutional guidelines with the approval of the Animal Care and Use Committee of the Tokyo University of Science.

### Tissue collection

Intravascular leukocytes were stained by i.v. injection of FITC-conjugated mAb (3 μg/mouse) against CD45.2 three minutes before sacrifice ^47^. Tumors and dLNs (right brachial lymph node) were collected.

### Preparing single cell suspension from dLN

dLN cells were prepared by standard method except for Fig. 5, in which dLN was digested for 45 minutes at 37°C with 0.1% collagenase (FUJIFILM Wako) then subjected to 25% Percoll PLUS (Cyvita) gradient and leukocytes were recovered from the pellet. Lineage (CD11b, B220, NK1.1, Ter119) positive cells were depleted using BD IMag Streptavidin Particles Plus-DM (BD Biosciences).

The cell concentration of the suspensions was determined using Flow-Count fluorospheres (Beckman Coulter) and a Cytoflex flow cytometer (Beckman Coulter). Cells were then stained with a mix of Fc Block (anti-mouse CD16/CD32 mAb; clone 2.4G2, BioLegend) and fluorophore-conjugated anti-mouse mAbs.

### Preparing single cell suspension from tumor

Tumors were cut into small fragments and digested for 45 minutes at 37°C with 0.1% collagenase (FUJIFILM Wako) in Fig. 1,2,4,5, and 0.1mg/mL Liberase TM (Roche) in Fig. 3. The cells were then subjected to 40% Percoll PLUS (Cytiva) and Histopaque-1083 (Sigma-Aldrich) density separation and leukocytes were recovered from the interphase. The cell concentration of the suspensions was determined using Flow-Count fluorospheres (Beckman) and a Cytoflex flow cytometer (Beckman). Cells were then stained with a mix of Fc Block and fluorophore-conjugated anti-mouse mAbs. TCRβ^+^ T cells were enriched magnetically using Anti-APC MicroBeads (Miltenyi) and LS columns (Miltenyi) except for single-cell RNA/TCRseq experiment (Fig. 3). In single-cell RNA/TCRseq experiment, cells were stained with Sample Tag oligonucleotide-conjugated antibodies in Single-Cell Multiplexing Kit (BD Biosciences) as in Supplementary Table S6.

### Flow cytometry and cell sorting

CD8^+^ (Fig. 1, 2, 3, 5), CD8^+^ PD1^−^, CD8^+^ PD1^+^ Ly108^+^ Tim3^−^, CD8^+^ PD1^+^ Ly108^+^ Tim3^+^, and CD8^+^ PD1^+^ Ly108^−^ Tim3^+^ T cells (Fig. 4) from tumor, and CD8^+^ CD44^high^ T cells from dLN were sorted using FACS Aria II or Aria III (BD Biosciences). Nonviable cells were excluded from the analysis based on forward and side scatter profiles and propidium iodide staining. Intravascular leukocytes were also excluded. Purity of sorted cells was always over 95%. Data were analyzed using FlowJo software (version 10.8.1; BD Biosciences).

### TCR library construction and sequencing

TCR sequencing libraries for next-generation sequencing were prepared according to the previous report ^27^. PolyA RNAs were isolated and amplified from sorted T cells according to a previous report (GSE110711; Shichino et al., 2019). To amplify the TCR cDNA containing complementarity determining region 3 (CDR3), nested PCR of the TCR locus was performed as follows. Beads containing cDNA was resuspended with the 12.5 μL of first PCR mixture comprised 0.5 μL of 10 μM trP1 primer, and 1 μL of TCR primer mix (Trac_ex, and Trbc_ex), 4.75 μL of water, and 6.25 μL of KAPA Hifi Hotstart ReadyMix (KAPA Biosystems). The thermal cycling conditions were programmed as follows: denaturation at 95°C for 3 minutes, 5 cycles of denaturation for 20 seconds at 98°C, annealing for 15 seconds at 65°C and extension for 30 seconds at 72°C, followed by a final extension at 72°C for 2 minutes. Next, 2.5 μL of the first-PCR products were added into the 22.5 μL of second PCR mixture consisted of 1.75 μL of 10 μM primer mix (trP1, Trac_in-Bio, and Trbc_in-Bio), 8.25 μL of water and 12.5 μL of KAPA Hifi Hotstart ReadyMix. The thermal cycling conditions were the same as the first PCR except for the cycle number, which was 25 cycles. Next, 25 μL of the second-PCR products were purified using AMPure XP (Beckman-Coulter) at a 0.7:1 ratio (beads-to-sample) and eluted in 20 μL of 10mM of Tris HCl pH 8.0.

Fragmentasion and adaptor ligation were performed using 10-20ng of second-PCR product as a template with NEBNext Ultra II FS DNA Library Prep Kit for Illumina (New England Biolabs) with some modifications: NEBNext Adaptor for Illumina in “Adaptor Ligation” step was substituted by 5uM P1 Adaptor (duplexed), and the reaction volume was 1/4 of the recommended in all steps. Adaptor-ligated DNA was purified using AM Pure XP kit (Beckman-Coulter) at a 0.8:1 ratio (beads to sample) and eluted in 20 μL of 10mM of Tris HCl pH 8.0. The third PCR was carried out using barcoded primers to enrich the TCRβ cDNA flanked with sequencing adapters. The third PCR mixture consisted of 1 μL of 10 μM trP1 primer, 1 μL of 10 μM IonA-BC[X]-Trbc primer, 3 μL of template and 5 μL of NEBNext Ultra II Q5 Master Mix (accessory of NEBNext Ultra II FS DNA Library Prep Kit). The thermal cycling conditions were programmed as follows: denaturation at 98°C for 30 seconds, 10 cycles of denaturation for 10 seconds at 98°C, annealing and extension for 1minute 15 seconds at 65°C, followed by a final extension at 65°C for 5 minutes. The PCR product was purified and subjected to size selection using AMPure XP kit (Beckman-Coulter) at a 1.0:1 ratio (beads to sample) and eluted with 20 μL of Tris-HCl (pH 8.0). Amplified TCRβ libraries were quantified using a KAPA SYBR Fast qPCR Kit (KAPA Biosystems) and size distribution was analyzed by Microchip Electrophoresis System MultiNA (Shimadzu).

Final TCRβ libraries, whose lengths were 200–300 base pairs, were pooled and sequenced using an Ion 540 Kit Chef, an Ion 540 Chip kit, and an Ion Genestudio S5 Sequencer (Thermo Fisher Scientific) according to the manufacturer’s instructions, except the flow number (500).

TCR sequencing libraries for next-generation sequencing were prepared according to the previous report ^44^. In brief, mRNA in T-cell lysate was captured by 10 μL of Dynabeads M270 streptavidin (Thermo Fisher Scientific) bound to 2.5 pmol of BioEcoP-dT25-adapter primers. To perform reverse transcription and template-switching, mRNA-trapped beads were suspended in 10 µL of RT mix [1× First Strand buffer (Thermo Fisher Scientific), 1 mM dNTP (Roche), 2.5 mM DTT (Thermo Fisher Scientific), 1 M betaine (Sigma-Aldrich), 9 mM MgCl2 (NIPPON GENE), 1 U/µL RNaseIn Plus RNase Inhibitor (Promega), 10 U/µL Superscript II (Thermo Fisher Scientific), and 1 µM of i5-TSO], and incubated for 60 min at 42°C and immediately cooled on ice. To amplify the TCR cDNA containing complementarity determining region 3 (CDR3), nested PCR of the TCR locus was performed following the purification of PCR product by an AM Pure XP kit (Beckman Coulter) at a 0.7:1 ratio of beads to sample and eluted with 20 μL of 10 mM Tris-HCl (pH 8.0). To amplify TCR libraries and add adaptor sequences for the next-generation sequencer, the third PCR was performed. The third-PCR products were purified as second PCR. The products were pooled and then purified and subjected to dual size selection using ProNex size-selective purification system (Promega) and eluted with 25 μL of 10 mM Tris-HCl (pH 8.5). Final TCR libraries, whose lengths were about 600 base pairs were sequenced using an Illumina Novaseq 6000 S4 flowcell (67 bp read 1 and 140 bp read 2) (Illumina). Only read2 contained the sequence regarding the definition of T cell clones.

### Library preparation and sequencing for scRNA/TCRseq

T cells from the spleen of Pmel Tg mouse, which stained with Single-Cell Multiplexing Kit (BD Biosciences) as in Supplementary Table S6, were spiked in after cell sorting. Live cells were stained with Calcein AM (Nacalai) and counted using Flow-Count fluorospheres (Beckman) and a Cytoflex flow cytometer (Beckman). About 18,000 labeled cells were trapped and reverse-transcribed using the BD Rhapsody™ (BD), Targeted mRNA and AbSeq Amplification Kit (BD), and Immune Response Panel Mm (BD) according to the manufacturer’s instructions. Targeted mRNA, TCRseq and Sample Tag libraries were prepared using according to the manufacturer’s instructions (VDJ CDR3 and Sample Tag Library Preparation Protocol) with the following modifications; (1) BD Rhapsody™ Immune Response Panel Mm with supplement panel including Cmpk2, Ifi204, Ifi211, Ifit1, Ifit3, Ifit3b, Ifit3b, Isg15, Mx1, Rsad2, and Usp18 were used for target mRNA amplification. (2) we used original set of primer for TCR amplification as summarized in Key resources table, and performed additional PCR (3^rd^ PCR, 6 cycles) to introduce unique dual index for sequencing. (3) we used ProNex Size-Selection DNA purification System (Promega) for purification of PCR product. Primers used for each PCR reaction and sample-to-beads ratios for purifying PCR product were summarized in Supplementary Table S7. Amplified libraries were quantified using a KAPA SYBR Fast qPCR Kit (KAPA Biosystems) and size distribution was analyzed by Microchip Electrophoresis System MultiNA (Shimadzu). Targeted RNAseq and Sample Tag libraries were sequenced on an Illumina Novaseq 6000 S4 flowcell (67 bp read 1 and 140 bp read 2) (Illumina) to a depth of approximately 40,000 reads and 10,000 reads per cell, respectively. TCRseq libraries were sequenced on an Illumina Novaseq 6000 SP flowcell (67 bp read 1 and 459 bp read 2) (Illumina) to a depth of approximately 20,000 reads.

### Data processing of TCR sequencing

Adapter trimming and quality filtering of sequencing data were performed by using Cutadapt-3.2 (Martin, 2013) and PRINSEQ-0.20.4 ^50^. Sequencing data were processed by MiXCR-3.0.5 ^51^. In MiXCR, filtered reads were aligned to reference mouse TCR V/D/J sequences registered in the international ImMunoGeneTics (IMGT) information system: -vParameters.geneFeatureToAlign = VTranscript - vjAlignmentOrder = JThenV, then identical sequences were assembled and grouped in clones with PCR and sequencing error correlation with the following parameters: - badQualityThreshold=10, –separateByV=true, --only-productive=true, –region-of-interest=CDR3. The Variable (V) and Joining (J) segment of TCRs were represented in IMGT gene nomenclature.

List of final clones were analyzed by VDJtools-1.2.1 ^52^. Then, the sequencing reads of sample was normalized to the cell count in each sample by “DownSample” command of VDJtools. T-cell clones were determined as TCR reads with the same TCR V segment, J segment and CDR3 nucleotide sequence.

### Calculation of clonality of TCR repertoire

The 1 - Pielou index was used to evaluate the clonality of TCR repertoire, which was calculated using the formula 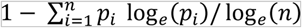, where *p_i_* is the frequency of clone *i* for a sample with *n* unique clones.

### Identification of OL clones in oligoclonal and polyclonal fraction

dLN-tumor OL clones were identified by “JoinSamples” function in VDJtools. These OL clones were ranked in order of their frequency in the tumor. OL clones that ranked in top 10 were defined as oligoclonal fraction of OL repertoire, and the other OL clones were defined as polyclonal fraction. In the scRNA/TCRseq analysis, OL clones were defined as those whose TCRb sequence (V segment, J segment and CDR3 nucleotide) was identified in the repertoire of dLN CD8^+^ CD44^high^ T cells, the output of bulk TCRseq. The oligoclonal/polyclona fraction in scTCR analysis on TILs was defined based on the frequency calculated in scTCRseq analysis.

### Data processing of scRNAseq reads

Data processing of scRNAseq reads was performed as described in Shichino et al. (TAS-seq). Adapter trimming of sequencing data was performed using cutadapt 2.10 (Martin, 2013). Filtered cell barcode reads were annotated by Python script provided by BD Biosciences with minor modification for compatibility to Python 3.7. Associated cDNA reads were mapped to reference RNA using bowtie2-2.4.2 ^53^ by the following parameters: -p 2 --very-sensitive-local -N 1 -norc -seed 656565 -reorder. Then, cell barcode information of each read was added to the bowtie2-mapped BAM files by the python script and pysam 0.15.4 ^54^, and read counts of each gene in each cell barcode were counted using mawk. Resultant count data was converted to a single-cell gene-expression matrix file by using R-3.6.3. The inflection point of the knee-plot (total read count versus the rank of the read count) was detected using DropletUtils package (v1.14.1, Lun *et al.*, 2018) in R-3.6.3. Cells of which total read count was over inflection point were considered as valid cells.

### Data processing of sample tag reads

For sample tag data, adapter trimming of sequencing data was performed by using cutadapt 3.4 (Martin, 2013). Filtered reads were chunked to 64 parts for parallel processing by using Seqkit 0.15.0 ^56^. Filtered cell barcode reads were annotated by Python script provided by BD with minor modification for compatible to Python3.8. Associated sample tag reads were mapped to known barcode fasta by using bowtie2-2.4.2 ^53^ by the following parameters: -p 2 -D 50 -R 20 -N 0 -L 14 -i S,1,0.75 –norc –seed 656565 – reorder –trim-to 3:40 –score-min L,-9,0 –mp 3,3 –np 3 –rdg 3,3.

Then, cell barcode information of each read were added to the bowtie2-mapped BAM files, and read counts of each Tag in each cell barcode were counted by using mawk. Resulted count data was converted to Tags x cells matrix file by using data.table package in R 3.6.3, and top 1M cell barcodes were extracted. For assignment of each tags to each cell barcodes, read counts of each tag in each valid cell barcode, which defined by the cDNA matrix, were extracted from tag/cell barcode expression matrix.

Unassigned cell barcodes were labeled as “not-detected” cells. Then, sum of the total read counts of each tags were normalized to the minimum sum count of each tags, and log_2_ fold-change between first most tag counts and second most tag counts within each cell barcode. Each cell barcode was ranked by the fold-change ascending order, and top N cells were identified as doublets (N was calculated as theoretically detectable doublets calculated by the Poisson’s distribution based on the number of loaded cells, total Rhapsody well number, and number of tags used).

### scRNAseq data analysis

scRNAseq data analysis was performed by using the R software package Seurat v4.0.1 ^57^. Expression data of targeted panel genes were converted to the Seurat object. Then, genes not included in the target panel and doublet or spiked-in Pmel cells were excluded. Expression data was log normalized and scaled using NormalizeData and ScaleData function, respectively. Principal component analysis (PCA) against all genes included in target panel was performed using RunPCA (number of calculated principal components (PCs) were 100), and enrichment of each PC was calculated using the JackStraw and ScoreJackStraw function (num.replicate = 100), and PCs that were significantly enriched (p ≤ 0.05) were selected for clustering and dimensional reduction analysis. Dimensional reduction was performed using RunUMAP. Cell clustering was performed using FindNeighbors and FindClusters against the significant PCs. Marker genes of each identified cluster were identified using FindAllMarkers function (test.use=“wilcox”, min.pct=0.1, return.thresh=0.05). Cluster constituted with contaminant cells determined by marker genes were excluded and the above analysis procedure was repeated.

### Transcriptional signature analysis in scRNAseq

We computed the extent to which gene signatures were expressed in cells using Seurat’s “AddModuleScore” function. Progenitor and terminally exhausted signature genes were obtained from the dataset of Miller et al. ^29^. Cytotoxicity and proliferation signature genes were selected according to the descriptions in BD Rhapsody™ Immune Response Panel Mm (BD). Genes used for calculating signature scores were summarized in Supplementary Table S8.

### Pseudotime analysis on scRNAseq data

Pseudotime analysis was performed using the Bioconductor package Slingshot v2.2.0 (Street et al., 2018). We ran the analysis using significantly enriched PCs (p ≤ 0.05) after setting Cluster 7 (NV-like) as the start cluster.

### Data processing of scTCRseq

Raw data of single-cell TCR sequencing were processed by pipeline produced by ImmunoGeneTeqs, Inc.. Briefly, sequencing reads were first separated by cell barcode, then adapter trimming and quality filtering of sequencing data were performed by using Cutadapt (Martin, 2013). PCR and sequencing error were corrected by lighter v1.1.2 ^59^ with following parameters: -newQual 25 -maxcor 4 -K 20. Error-corrected sequence reads were processed by MiXCR ^51^ to generate the list of TCRα or TCRβ sequences for each cell barcode, using reference mouse TCR V/D/J sequences registered in the international IMGT information system with the parameters summarized in Supplementary Table S9. Quality filtering of cell barcodes was performed with the following criteria: (1) more than 32 TCR reads were detected. (2) the proportion of the most frequent TCR sequence in cell barcode was over 0.6. (3) the cell barcode was annotated to the T cell cluster in scRNAseq analysis. Finally, the most frequent sequence of TCRα and TCRβ was adopted and paired in each cell barcode. Identified TCRα/β sequence was imported into the meta.data matrix in Seurat object of scRNAseq data based on the cellular barcode identities. T-cell clones were determined as cells with the same TCRα and TCRβ pair, defined by V segment, J segment and CDR3 nucleotide sequence. Frequency of each clone was calculated as the proportion of cells belonging to the clone in all T cells whose TCRα and TCRβ pair was assigned.

### Evaluating the exhaustion status of T cell clones by TCRseq on CD8^+^ TIL subsets

To evaluate the exhaustion status of OL clones in oligoclonal and polyclonal fraction, we first calculated the frequency of clones in “Pooled” CD8^+^ TIL repertoire from the repertoire of TIL subsets (PD1^−^, Ly108, DP, and Tim-3) as the following equation:

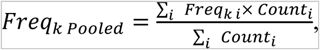

where *Freq* means the frequency of each clone k, and *Count* means the cell count of each subset i. Then, OL clones in oligoclonal and polyclonal fraction was identified based on their frequency in Pooled repertoire as above. The total frequency of these clones in each TIL subset was calculated, and defined as oligoclonal and polyclonal OL repertoire in each TIL subset.

### Quantification and statistical analysis

Statistical analyses were performed using GraphPad Prism (ver9.2.0) software (GraphPad Software) except for correlation analysis in Fig. 2e and 3g. One-way ANOVA with Tukey’s post test was run on the comparison between treatment groups. Spearman correlation test was run on the correlation analysis using cor.test function in Microsoft R open 4.1.1. One-way Repeated-Measures (RM) ANOVA with Tukey’s post test was run on the comparison between CD8^+^ TIL subsets. Paired t test was run on the comparison between the dominance of oligoclonal or polyclonal fraction of dLN-tumor OL. Multiple unpaired t tests was run on the comparison between DT treated and untreated mice for each treatment group. Asterisks to indicate significance corresponding to the following: n.s., not significant (*P*>0.05), \**P* ≤ 0.05, \*\**P* ≤ 0.01, *** *P* ≤ 0.001.

## Acknowledgements

We would like to thank Y. Hara for advice in cell sorting, staff of RIBS animal facility for supporting the maintenance of animals, member of IGT. Inc. for expert technical assistance in TCR sequencing, D. Komura and S. Ishikawa for advice in data analysis. We would also like to thank Editage (www.editage.com) for English language editing. Fig. 5a was created with BioRender.com.

This work was supported by the Japan Society for the Promotion of Science under Grant Number 20281832 and 17929397, and by the Japan Agency for Medical Research and Development (AMED) under Grant Number JP 21gm6210025, 22fk0310509s0101, and 22ama221306h0001.

## Author Contributions

H.A., K.M., and S.U. designed research. H.A. and S.U. performed research. T.A. generated mice. H.A., M.T., H.O., H.S., H.A., and S.U. processed the samples. H.A. analyzed data with supports from S.S. and S.U.. H.A. and S.U. wrote the initial draft of the manuscript. All the authors participated in writing the final manuscript.

