## Supplemental Figures and Tables for "Clonal spreading of tumor-infiltrating T cells underlies the robust antitumor immune responses"

### Slide 1
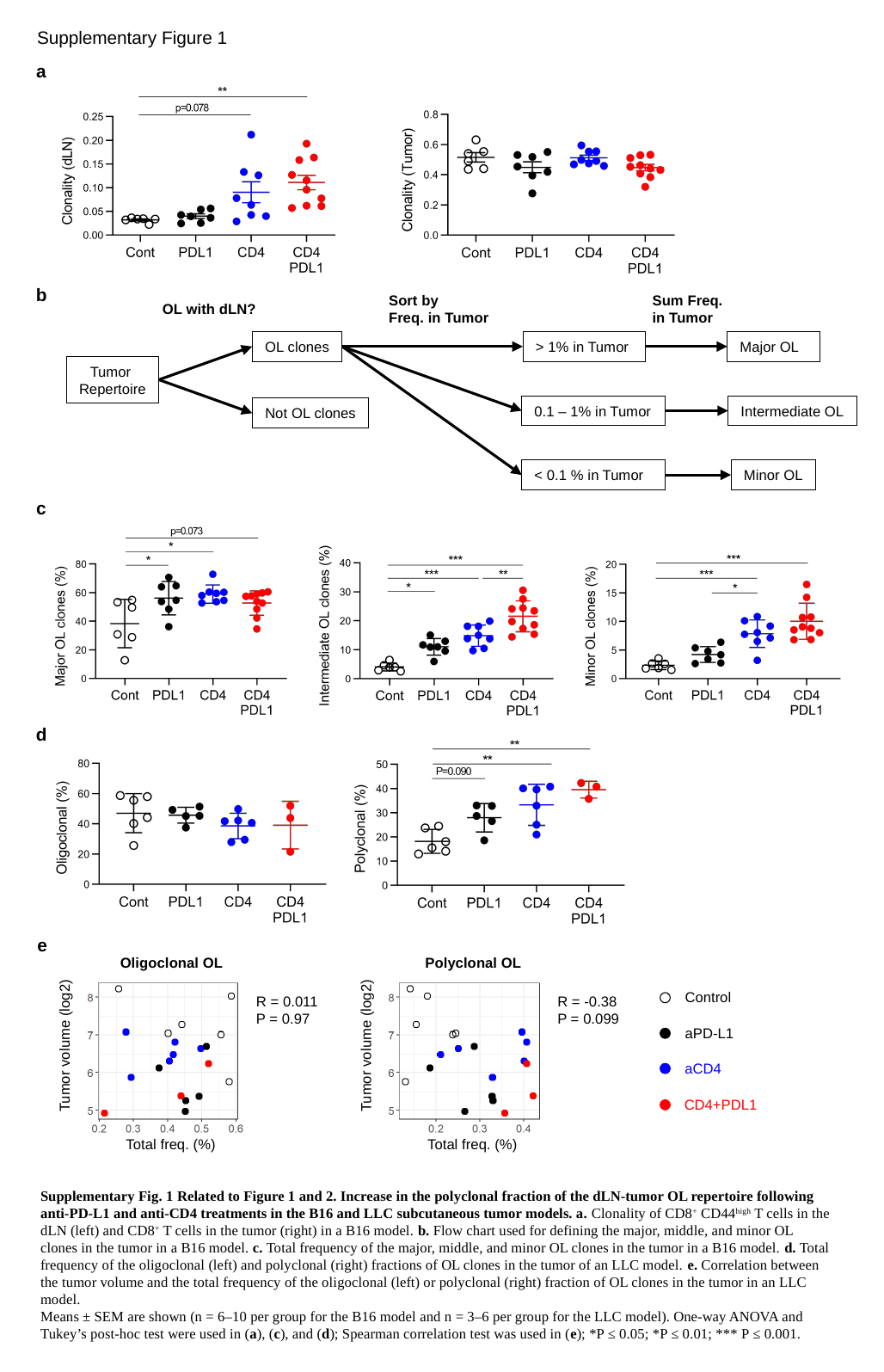

Supplementary Figure 1
a
b
Sort by
Freq. in Tumor
Sum Freq.
in Tumor
OL with dLN?
OL clones
> 1% in Tumor
Major OL
Tumor
Repertoire
0.1 – 1% in Tumor
Intermediate OL
Not OL clones
< 0.1 % in Tumor
Minor OL
c
d
e
Oligoclonal OL
Polyclonal OL
Control
aPD-L1
aCD4
CD4+PDL1
R = 0.011
P = 0.97
R = -0.38
P = 0.099
Tumor volume (log2)
Tumor volume (log2)
Total freq. (%)
Total freq. (%)
Supplementary Fig. 1 Related to Figure 1 and 2. Increase in the polyclonal fraction of the dLN-tumor OL repertoire following anti-PD-L1 and anti-CD4 treatments in the B16 and LLC subcutaneous tumor models. a. Clonality of CD8^+^ CD44^high^ T cells in the dLN (left) and CD8^+^ T cells in the tumor (right) in a B16 model. b. Flow chart used for defining the major, middle, and minor OL clones in the tumor in a B16 model. c. Total frequency of the major, middle, and minor OL clones in the tumor in a B16 model. d. Total frequency of the oligoclonal (left) and polyclonal (right) fractions of OL clones in the tumor of an LLC model. e. Correlation between the tumor volume and the total frequency of the oligoclonal (left) or polyclonal (right) fraction of OL clones in the tumor in an LLC model.
Means ± SEM are shown (n = 6–10 per group for the B16 model and n = 3–6 per group for the LLC model). One-way ANOVA and Tukey’s post-hoc test were used in (a), (c), and (d); Spearman correlation test was used in (e); *P ≤ 0.05; *P ≤ 0.01; *** P ≤ 0.001.

### Slide 2
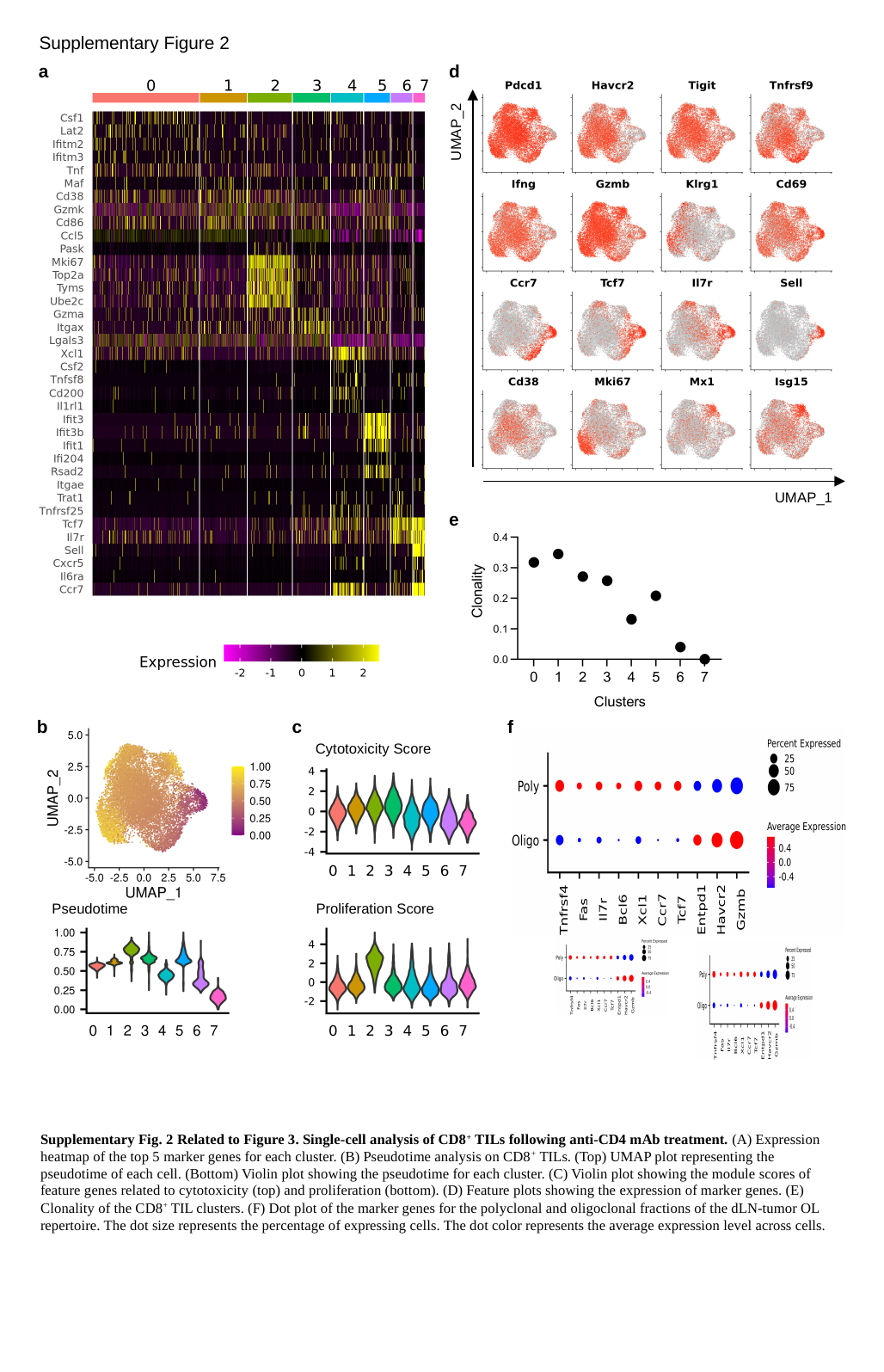

Supplementary Figure 2
a
d
UMAP_2
UMAP_1
e
b
Pseudotime
c
Cytotoxicity Score
Proliferation Score
f
Supplementary Fig. 2 Related to Figure 3. Single-cell analysis of CD8^+^ TILs following anti-CD4 mAb treatment. (A) Expression heatmap of the top 5 marker genes for each cluster. (B) Pseudotime analysis on CD8^+^ TILs. (Top) UMAP plot representing the pseudotime of each cell. (Bottom) Violin plot showing the pseudotime for each cluster. (C) Violin plot showing the module scores of feature genes related to cytotoxicity (top) and proliferation (bottom). (D) Feature plots showing the expression of marker genes. (E) Clonality of the CD8^+^ TIL clusters. (F) Dot plot of the marker genes for the polyclonal and oligoclonal fractions of the dLN-tumor OL repertoire. The dot size represents the percentage of expressing cells. The dot color represents the average expression level across cells.

### Slide 3
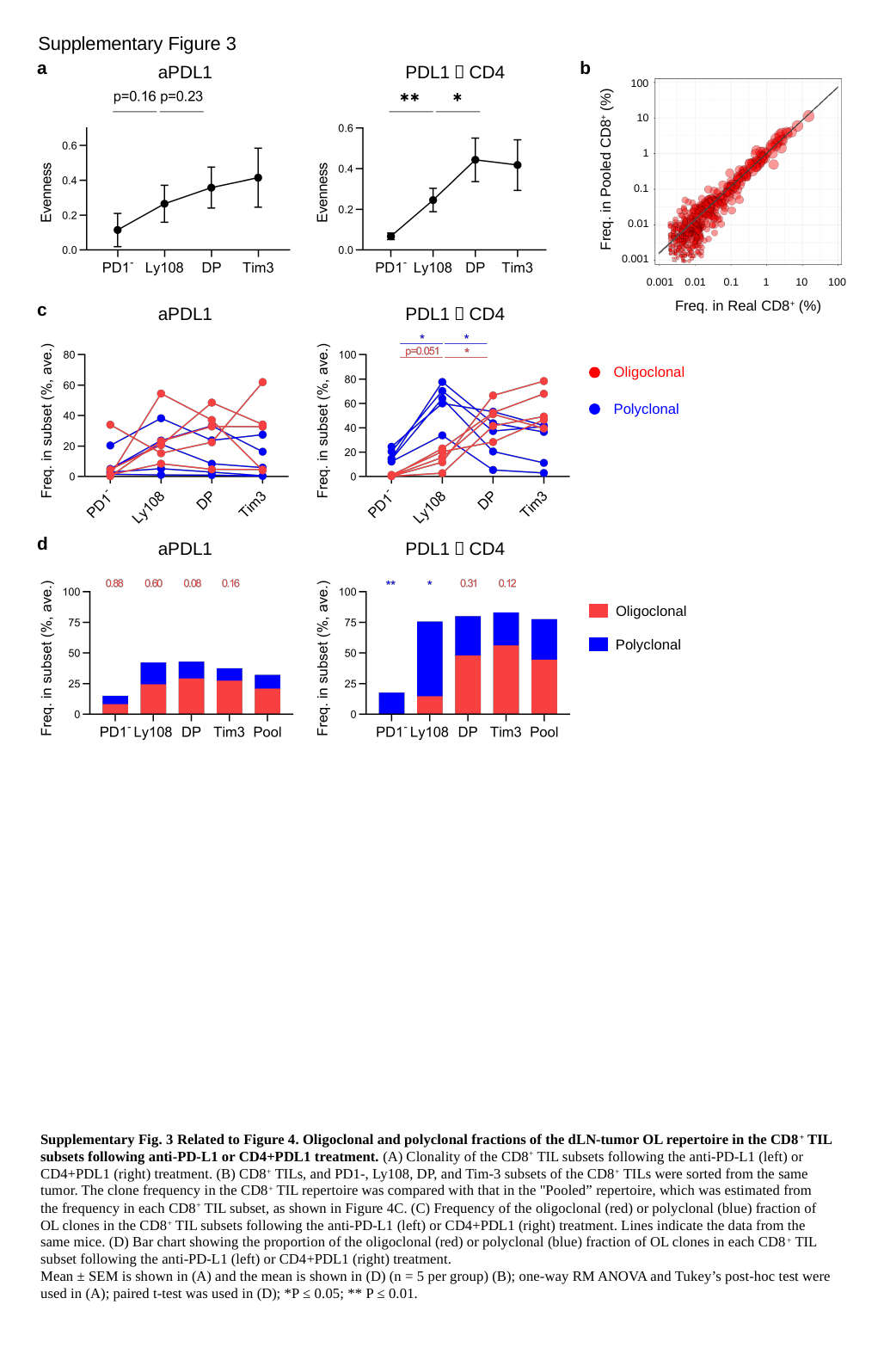

Supplementary Figure 3
a
b
aPDL1
PDL1＋CD4
100
10
1
Freq. in Pooled CD8^+^ (%)
0.1
0.01
0.001
0.001
0.01
0.1
1
10
100
Freq. in Real CD8^+^ (%)
c
aPDL1
PDL1＋CD4
Oligoclonal
Polyclonal
d
aPDL1
PDL1＋CD4
Oligoclonal
Polyclonal
Supplementary Fig. 3 Related to Figure 4. Oligoclonal and polyclonal fractions of the dLN-tumor OL repertoire in the CD8^+^ TIL subsets following anti-PD-L1 or CD4+PDL1 treatment. (A) Clonality of the CD8^+^ TIL subsets following the anti-PD-L1 (left) or CD4+PDL1 (right) treatment. (B) CD8^+^ TILs, and PD1-, Ly108, DP, and Tim-3 subsets of the CD8^+^ TILs were sorted from the same tumor. The clone frequency in the CD8^+^ TIL repertoire was compared with that in the "Pooled” repertoire, which was estimated from the frequency in each CD8^+^ TIL subset, as shown in Figure 4C. (C) Frequency of the oligoclonal (red) or polyclonal (blue) fraction of OL clones in the CD8^+^ TIL subsets following the anti-PD-L1 (left) or CD4+PDL1 (right) treatment. Lines indicate the data from the same mice. (D) Bar chart showing the proportion of the oligoclonal (red) or polyclonal (blue) fraction of OL clones in each CD8^+^ TIL subset following the anti-PD-L1 (left) or CD4+PDL1 (right) treatment.
Mean ± SEM is shown in (A) and the mean is shown in (D) (n = 5 per group) (B); one-way RM ANOVA and Tukey’s post-hoc test were used in (A); paired t-test was used in (D); *P ≤ 0.05; ** P ≤ 0.01.

### Slide 4
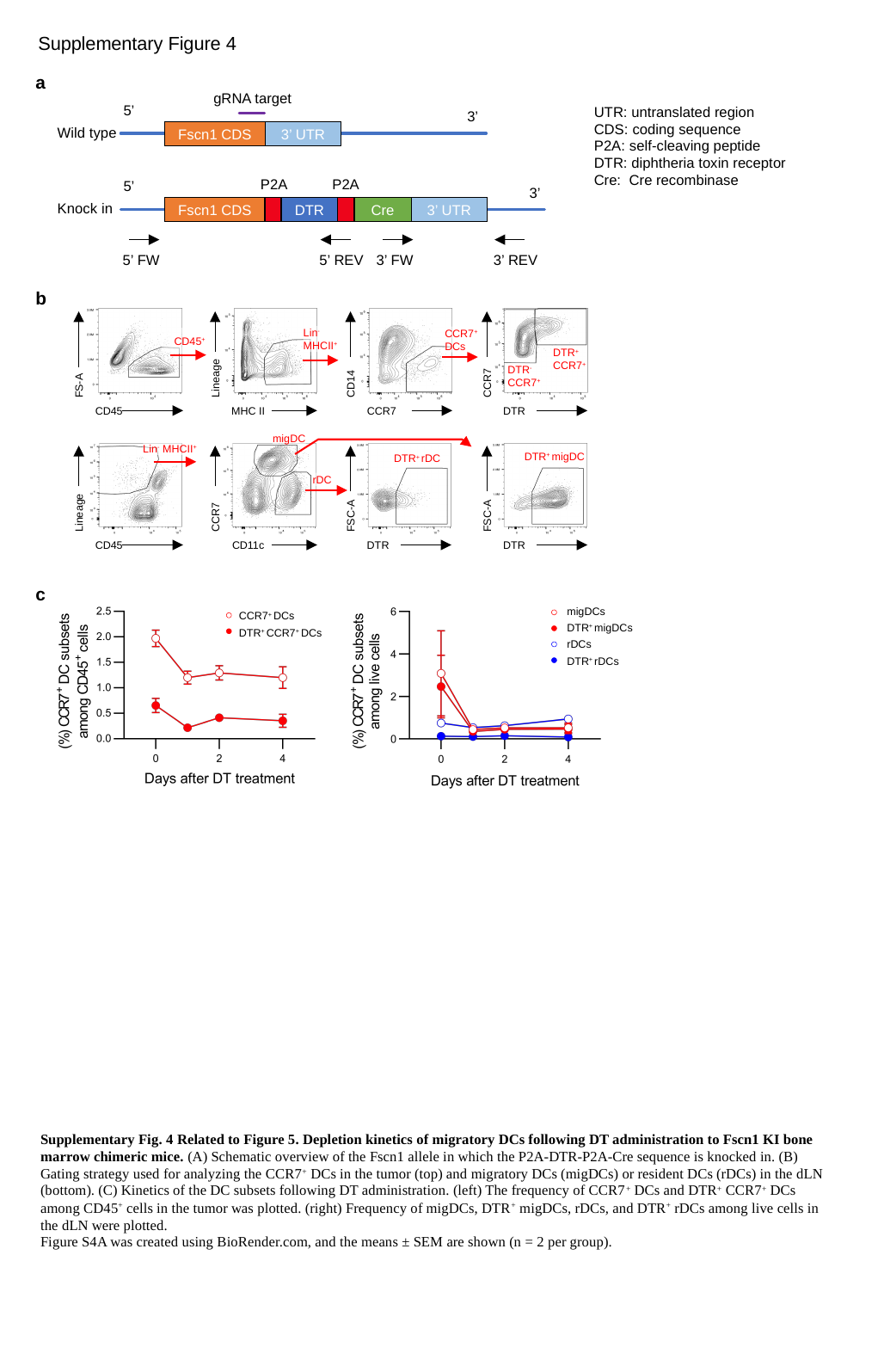

Supplementary Figure 4
a
gRNA target
5’
UTR: untranslated region
CDS: coding sequence
P2A: self-cleaving peptide
DTR: diphtheria toxin receptor
Cre: Cre recombinase
3’
Wild type
Fscn1 CDS
3’ UTR
P2A
P2A
5’
3’
Knock in
Fscn1 CDS
DTR
Cre
3’ UTR
5’ FW
5’ REV
3’ FW
3’ REV
b
Lin^-^
MHCII^+^
CCR7^+^
DCs
CD45^+^
DTR^+^
CCR7^+^
DTR^-^
CCR7^+^
Lineage
CCR7
CD14
FS-A
CD45
MHC II
CCR7
DTR
migDC
Lin^-^ MHCII^+^
DTR^+^ migDC
DTR^+^ rDC
rDC
Lineage
FSC-A
FSC-A
CCR7
CD45
CD11c
DTR
DTR
c
migDCs
DTR^+^ migDCs
rDCs
DTR^+^ rDCs
CCR7^+^ DCs
DTR^+^ CCR7^+^ DCs
Supplementary Fig. 4 Related to Figure 5. Depletion kinetics of migratory DCs following DT administration to Fscn1 KI bone marrow chimeric mice. (A) Schematic overview of the Fscn1 allele in which the P2A-DTR-P2A-Cre sequence is knocked in. (B) Gating strategy used for analyzing the CCR7^+^ DCs in the tumor (top) and migratory DCs (migDCs) or resident DCs (rDCs) in the dLN (bottom). (C) Kinetics of the DC subsets following DT administration. (left) The frequency of CCR7^+^ DCs and DTR^+^ CCR7^+^ DCs among CD45^+^ cells in the tumor was plotted. (right) Frequency of migDCs, DTR^+^ migDCs, rDCs, and DTR^+^ rDCs among live cells in the dLN were plotted.
Figure S4A was created using BioRender.com, and the means ± SEM are shown (n = 2 per group).

### Slide 5
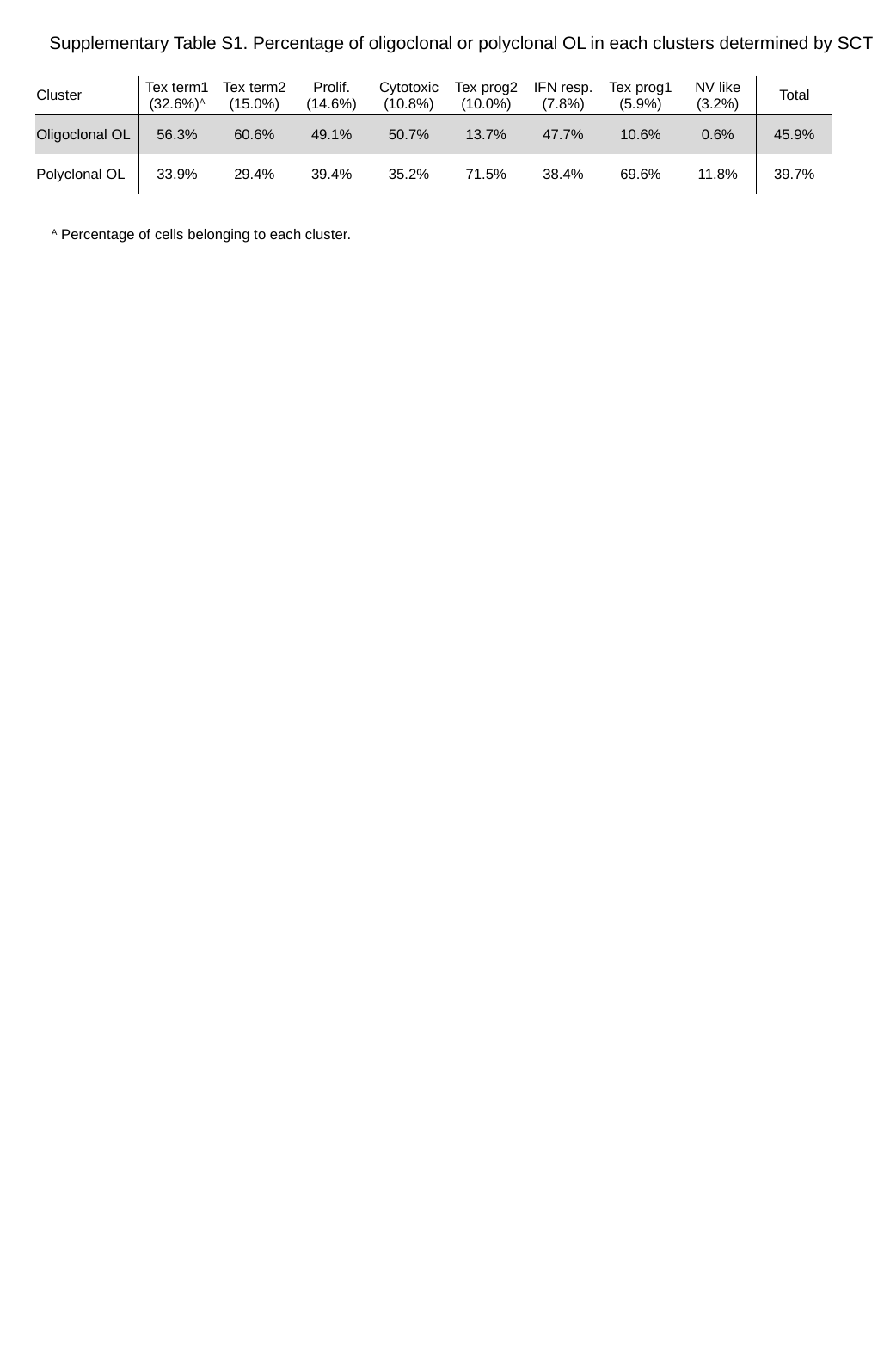

Supplementary Table S1. Percentage of oligoclonal or polyclonal OL in each clusters determined by SCT
| Cluster | Tex term1 (32.6%)^A^ | Tex term2 (15.0%) | Prolif. (14.6%) | Cytotoxic (10.8%) | Tex prog2 (10.0%) | IFN resp. (7.8%) | Tex prog1 (5.9%) | NV like (3.2%) | Total |
| --- | --- | --- | --- | --- | --- | --- | --- | --- | --- |
| Oligoclonal OL | 56.3% | 60.6% | 49.1% | 50.7% | 13.7% | 47.7% | 10.6% | 0.6% | 45.9% |
| Polyclonal OL | 33.9% | 29.4% | 39.4% | 35.2% | 71.5% | 38.4% | 69.6% | 11.8% | 39.7% |
^A^ Percentage of cells belonging to each cluster.

### Slide 6
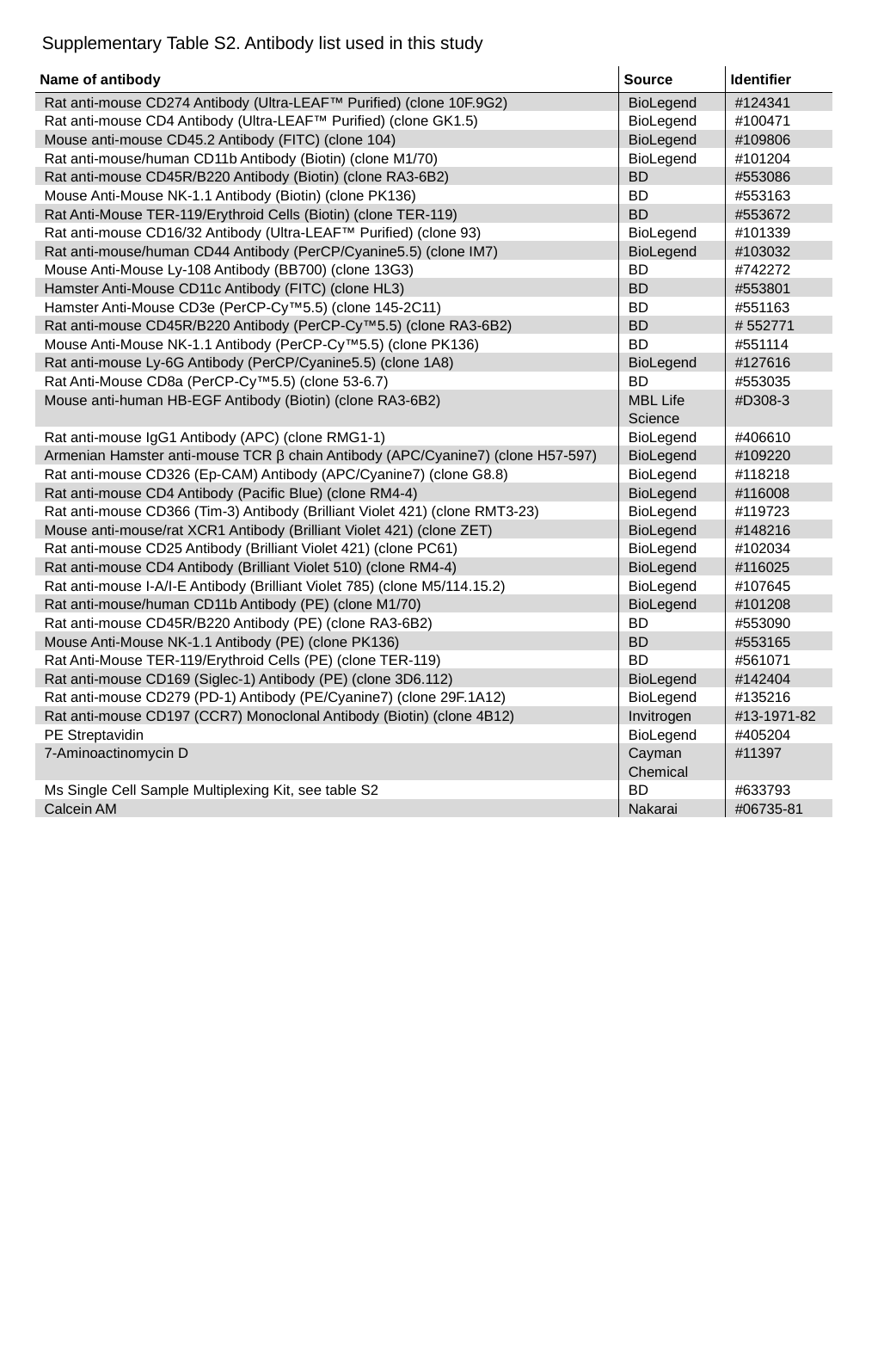

Supplementary Table S2. Antibody list used in this study
| Name of antibody | Source | Identifier |
| --- | --- | --- |
| Rat anti-mouse CD274 Antibody (Ultra-LEAF™ Purified) (clone 10F.9G2) | BioLegend | #124341 |
| Rat anti-mouse CD4 Antibody (Ultra-LEAF™ Purified) (clone GK1.5) | BioLegend | #100471 |
| Mouse anti-mouse CD45.2 Antibody (FITC) (clone 104) | BioLegend | #109806 |
| Rat anti-mouse/human CD11b Antibody (Biotin) (clone M1/70) | BioLegend | #101204 |
| Rat anti-mouse CD45R/B220 Antibody (Biotin) (clone RA3-6B2) | BD | #553086 |
| Mouse Anti-Mouse NK-1.1 Antibody (Biotin) (clone PK136) | BD | #553163 |
| Rat Anti-Mouse TER-119/Erythroid Cells (Biotin) (clone TER-119) | BD | #553672 |
| Rat anti-mouse CD16/32 Antibody (Ultra-LEAF™ Purified) (clone 93) | BioLegend | #101339 |
| Rat anti-mouse/human CD44 Antibody (PerCP/Cyanine5.5) (clone IM7) | BioLegend | #103032 |
| Mouse Anti-Mouse Ly-108 Antibody (BB700) (clone 13G3) | BD | #742272 |
| Hamster Anti-Mouse CD11c Antibody (FITC) (clone HL3) | BD | #553801 |
| Hamster Anti-Mouse CD3e (PerCP-Cy™5.5) (clone 145-2C11) | BD | #551163 |
| Rat anti-mouse CD45R/B220 Antibody (PerCP-Cy™5.5) (clone RA3-6B2) | BD | # 552771 |
| Mouse Anti-Mouse NK-1.1 Antibody (PerCP-Cy™5.5) (clone PK136) | BD | #551114 |
| Rat anti-mouse Ly-6G Antibody (PerCP/Cyanine5.5) (clone 1A8) | BioLegend | #127616 |
| Rat Anti-Mouse CD8a (PerCP-Cy™5.5) (clone 53-6.7) | BD | #553035 |
| Mouse anti-human HB-EGF Antibody (Biotin) (clone RA3-6B2) | MBL Life Science | #D308-3 |
| Rat anti-mouse IgG1 Antibody (APC) (clone RMG1-1) | BioLegend | #406610 |
| Armenian Hamster anti-mouse TCR β chain Antibody (APC/Cyanine7) (clone H57-597) | BioLegend | #109220 |
| Rat anti-mouse CD326 (Ep-CAM) Antibody (APC/Cyanine7) (clone G8.8) | BioLegend | #118218 |
| Rat anti-mouse CD4 Antibody (Pacific Blue) (clone RM4-4) | BioLegend | #116008 |
| Rat anti-mouse CD366 (Tim-3) Antibody (Brilliant Violet 421) (clone RMT3-23) | BioLegend | #119723 |
| Mouse anti-mouse/rat XCR1 Antibody (Brilliant Violet 421) (clone ZET) | BioLegend | #148216 |
| Rat anti-mouse CD25 Antibody (Brilliant Violet 421) (clone PC61) | BioLegend | #102034 |
| Rat anti-mouse CD4 Antibody (Brilliant Violet 510) (clone RM4-4) | BioLegend | #116025 |
| Rat anti-mouse I-A/I-E Antibody (Brilliant Violet 785) (clone M5/114.15.2) | BioLegend | #107645 |
| Rat anti-mouse/human CD11b Antibody (PE) (clone M1/70) | BioLegend | #101208 |
| Rat anti-mouse CD45R/B220 Antibody (PE) (clone RA3-6B2) | BD | #553090 |
| Mouse Anti-Mouse NK-1.1 Antibody (PE) (clone PK136) | BD | #553165 |
| Rat Anti-Mouse TER-119/Erythroid Cells (PE) (clone TER-119) | BD | #561071 |
| Rat anti-mouse CD169 (Siglec-1) Antibody (PE) (clone 3D6.112) | BioLegend | #142404 |
| Rat anti-mouse CD279 (PD-1) Antibody (PE/Cyanine7) (clone 29F.1A12) | BioLegend | #135216 |
| Rat anti-mouse CD197 (CCR7) Monoclonal Antibody (Biotin) (clone 4B12) | Invitrogen | #13-1971-82 |
| PE Streptavidin | BioLegend | #405204 |
| 7-Aminoactinomycin D | Cayman Chemical | #11397 |
| Ms Single Cell Sample Multiplexing Kit, see table S2 | BD | #633793 |
| Calcein AM | Nakarai | #06735-81 |

### Slide 7
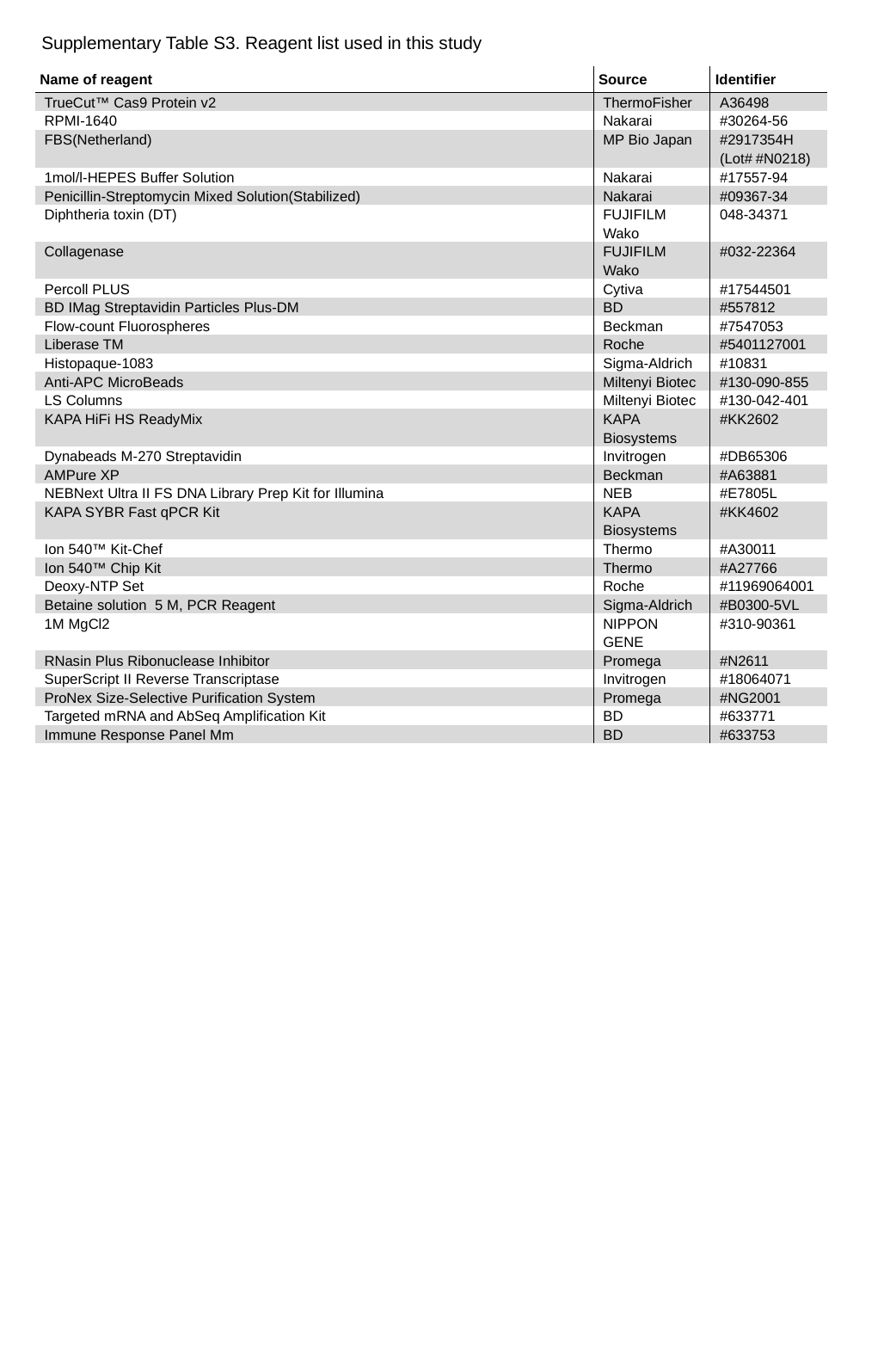

Supplementary Table S3. Reagent list used in this study
| Name of reagent | Source | Identifier |
| --- | --- | --- |
| TrueCut™ Cas9 Protein v2 | ThermoFisher | A36498 |
| RPMI-1640 | Nakarai | #30264-56 |
| FBS(Netherland) | MP Bio Japan | #2917354H (Lot# #N0218) |
| 1mol/l-HEPES Buffer Solution | Nakarai | #17557-94 |
| Penicillin-Streptomycin Mixed Solution(Stabilized) | Nakarai | #09367-34 |
| Diphtheria toxin (DT) | FUJIFILM Wako | 048-34371 |
| Collagenase | FUJIFILM Wako | #032-22364 |
| Percoll PLUS | Cytiva | #17544501 |
| BD IMag Streptavidin Particles Plus-DM | BD | #557812 |
| Flow-count Fluorospheres | Beckman | #7547053 |
| Liberase TM | Roche | #5401127001 |
| Histopaque-1083 | Sigma-Aldrich | #10831 |
| Anti-APC MicroBeads | Miltenyi Biotec | #130-090-855 |
| LS Columns | Miltenyi Biotec | #130-042-401 |
| KAPA HiFi HS ReadyMix | KAPA Biosystems | #KK2602 |
| Dynabeads M-270 Streptavidin | Invitrogen | #DB65306 |
| AMPure XP | Beckman | #A63881 |
| NEBNext Ultra II FS DNA Library Prep Kit for Illumina | NEB | #E7805L |
| KAPA SYBR Fast qPCR Kit | KAPA Biosystems | #KK4602 |
| Ion 540™ Kit-Chef | Thermo | #A30011 |
| Ion 540™ Chip Kit | Thermo | #A27766 |
| Deoxy-NTP Set | Roche | #11969064001 |
| Betaine solution 5 M, PCR Reagent | Sigma-Aldrich | #B0300-5VL |
| 1M MgCl2 | NIPPON GENE | #310-90361 |
| RNasin Plus Ribonuclease Inhibitor | Promega | #N2611 |
| SuperScript II Reverse Transcriptase | Invitrogen | #18064071 |
| ProNex Size-Selective Purification System | Promega | #NG2001 |
| Targeted mRNA and AbSeq Amplification Kit | BD | #633771 |
| Immune Response Panel Mm | BD | #633753 |

### Slide 8
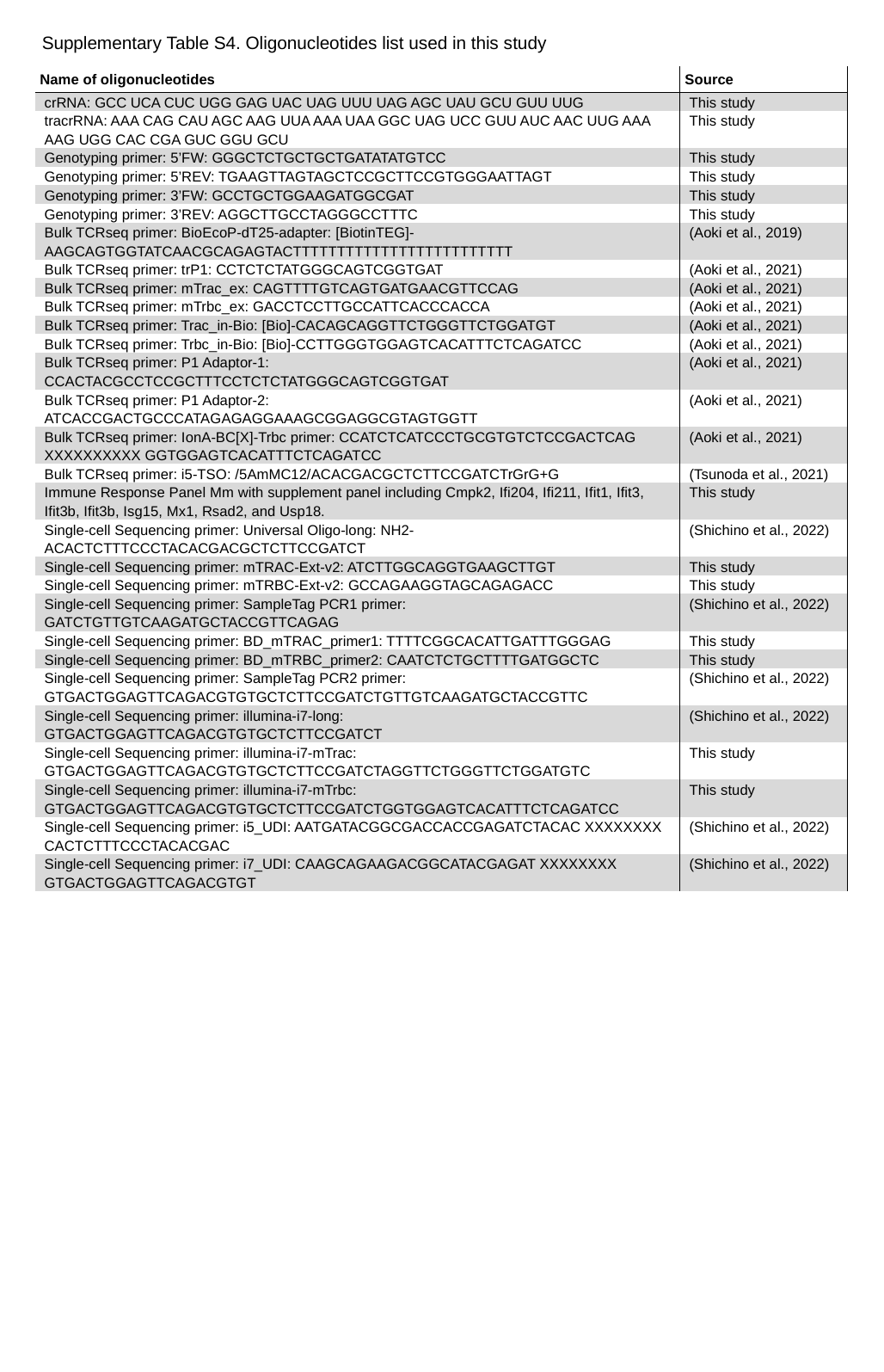

Supplementary Table S4. Oligonucleotides list used in this study
| Name of oligonucleotides | Source |
| --- | --- |
| crRNA: GCC UCA CUC UGG GAG UAC UAG UUU UAG AGC UAU GCU GUU UUG | This study |
| tracrRNA: AAA CAG CAU AGC AAG UUA AAA UAA GGC UAG UCC GUU AUC AAC UUG AAA AAG UGG CAC CGA GUC GGU GCU | This study |
| Genotyping primer: 5’FW: GGGCTCTGCTGCTGATATATGTCC | This study |
| Genotyping primer: 5’REV: TGAAGTTAGTAGCTCCGCTTCCGTGGGAATTAGT | This study |
| Genotyping primer: 3’FW: GCCTGCTGGAAGATGGCGAT | This study |
| Genotyping primer: 3’REV: AGGCTTGCCTAGGGCCTTTC | This study |
| Bulk TCRseq primer: BioEcoP-dT25-adapter: [BiotinTEG]-AAGCAGTGGTATCAACGCAGAGTACTTTTTTTTTTTTTTTTTTTTTTTTT | (Aoki et al., 2019) |
| Bulk TCRseq primer: trP1: CCTCTCTATGGGCAGTCGGTGAT | (Aoki et al., 2021) |
| Bulk TCRseq primer: mTrac\_ex: CAGTTTTGTCAGTGATGAACGTTCCAG | (Aoki et al., 2021) |
| Bulk TCRseq primer: mTrbc\_ex: GACCTCCTTGCCATTCACCCACCA | (Aoki et al., 2021) |
| Bulk TCRseq primer: Trac\_in-Bio: [Bio]-CACAGCAGGTTCTGGGTTCTGGATGT | (Aoki et al., 2021) |
| Bulk TCRseq primer: Trbc\_in-Bio: [Bio]-CCTTGGGTGGAGTCACATTTCTCAGATCC | (Aoki et al., 2021) |
| Bulk TCRseq primer: P1 Adaptor-1: CCACTACGCCTCCGCTTTCCTCTCTATGGGCAGTCGGTGAT | (Aoki et al., 2021) |
| Bulk TCRseq primer: P1 Adaptor-2: ATCACCGACTGCCCATAGAGAGGAAAGCGGAGGCGTAGTGGTT | (Aoki et al., 2021) |
| Bulk TCRseq primer: IonA-BC[X]-Trbc primer: CCATCTCATCCCTGCGTGTCTCCGACTCAG XXXXXXXXXX GGTGGAGTCACATTTCTCAGATCC | (Aoki et al., 2021) |
| Bulk TCRseq primer: i5-TSO: /5AmMC12/ACACGACGCTCTTCCGATCTrGrG+G | (Tsunoda et al., 2021) |
| Immune Response Panel Mm with supplement panel including Cmpk2, Ifi204, Ifi211, Ifit1, Ifit3, Ifit3b, Ifit3b, Isg15, Mx1, Rsad2, and Usp18. | This study |
| Single-cell Sequencing primer: Universal Oligo-long: NH2-ACACTCTTTCCCTACACGACGCTCTTCCGATCT | (Shichino et al., 2022) |
| Single-cell Sequencing primer: mTRAC-Ext-v2: ATCTTGGCAGGTGAAGCTTGT | This study |
| Single-cell Sequencing primer: mTRBC-Ext-v2: GCCAGAAGGTAGCAGAGACC | This study |
| Single-cell Sequencing primer: SampleTag PCR1 primer: GATCTGTTGTCAAGATGCTACCGTTCAGAG | (Shichino et al., 2022) |
| Single-cell Sequencing primer: BD\_mTRAC\_primer1: TTTTCGGCACATTGATTTGGGAG | This study |
| Single-cell Sequencing primer: BD\_mTRBC\_primer2: CAATCTCTGCTTTTGATGGCTC | This study |
| Single-cell Sequencing primer: SampleTag PCR2 primer: GTGACTGGAGTTCAGACGTGTGCTCTTCCGATCTGTTGTCAAGATGCTACCGTTC | (Shichino et al., 2022) |
| Single-cell Sequencing primer: illumina-i7-long: GTGACTGGAGTTCAGACGTGTGCTCTTCCGATCT | (Shichino et al., 2022) |
| Single-cell Sequencing primer: illumina-i7-mTrac: GTGACTGGAGTTCAGACGTGTGCTCTTCCGATCTAGGTTCTGGGTTCTGGATGTC | This study |
| Single-cell Sequencing primer: illumina-i7-mTrbc: GTGACTGGAGTTCAGACGTGTGCTCTTCCGATCTGGTGGAGTCACATTTCTCAGATCC | This study |
| Single-cell Sequencing primer: i5\_UDI: AATGATACGGCGACCACCGAGATCTACAC XXXXXXXX CACTCTTTCCCTACACGAC | (Shichino et al., 2022) |
| Single-cell Sequencing primer: i7\_UDI: CAAGCAGAAGACGGCATACGAGAT XXXXXXXX GTGACTGGAGTTCAGACGTGT | (Shichino et al., 2022) |

### Slide 9
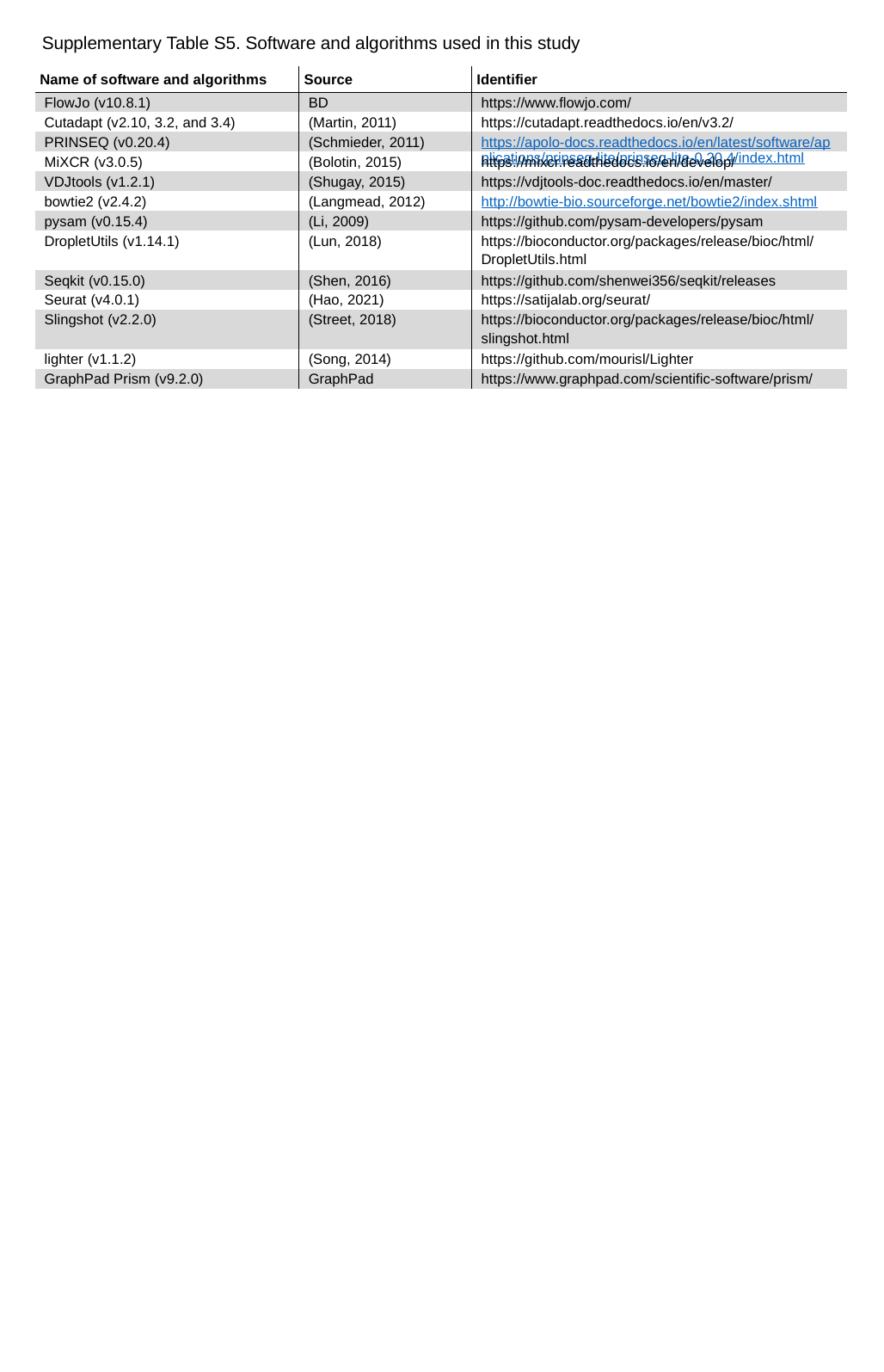

Supplementary Table S5. Software and algorithms used in this study
| Name of software and algorithms | Source | Identifier |
| --- | --- | --- |
| FlowJo (v10.8.1) | BD | https://www.flowjo.com/ |
| Cutadapt (v2.10, 3.2, and 3.4) | (Martin, 2011) | https://cutadapt.readthedocs.io/en/v3.2/ |
| PRINSEQ (v0.20.4) | (Schmieder, 2011) | https://apolo-docs.readthedocs.io/en/latest/software/applications/prinseq-lite/prinseq-lite-0.20.4/index.html |
| MiXCR (v3.0.5) | (Bolotin, 2015) | https://mixcr.readthedocs.io/en/develop/ |
| VDJtools (v1.2.1) | (Shugay, 2015) | https://vdjtools-doc.readthedocs.io/en/master/ |
| bowtie2 (v2.4.2) | (Langmead, 2012) | http://bowtie-bio.sourceforge.net/bowtie2/index.shtml |
| pysam (v0.15.4) | (Li, 2009) | https://github.com/pysam-developers/pysam |
| DropletUtils (v1.14.1) | (Lun, 2018) | https://bioconductor.org/packages/release/bioc/html/DropletUtils.html |
| Seqkit (v0.15.0) | (Shen, 2016) | https://github.com/shenwei356/seqkit/releases |
| Seurat (v4.0.1) | (Hao, 2021) | https://satijalab.org/seurat/ |
| Slingshot (v2.2.0) | (Street, 2018) | https://bioconductor.org/packages/release/bioc/html/slingshot.html |
| lighter (v1.1.2) | (Song, 2014) | https://github.com/mourisl/Lighter |
| GraphPad Prism (v9.2.0) | GraphPad | https://www.graphpad.com/scientific-software/prism/ |

### Slide 10
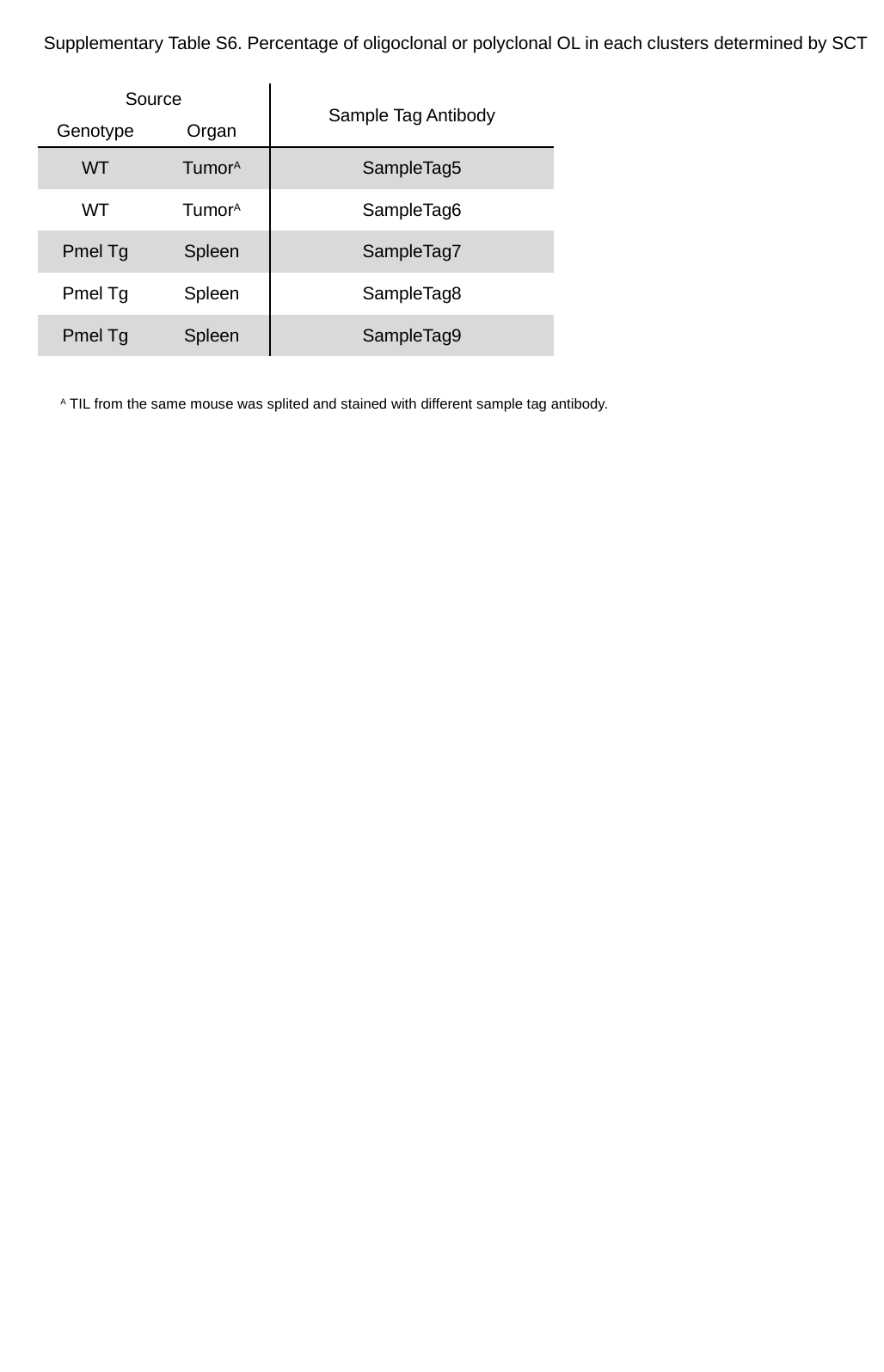

Supplementary Table S6. Percentage of oligoclonal or polyclonal OL in each clusters determined by SCT
| Source | | Sample Tag Antibody |
| --- | --- | --- |
| Genotype | Organ | |
| WT | Tumor^A^ | SampleTag5 |
| WT | Tumor^A^ | SampleTag6 |
| Pmel Tg | Spleen | SampleTag7 |
| Pmel Tg | Spleen | SampleTag8 |
| Pmel Tg | Spleen | SampleTag9 |
^A^ TIL from the same mouse was splited and stained with different sample tag antibody.

### Slide 11
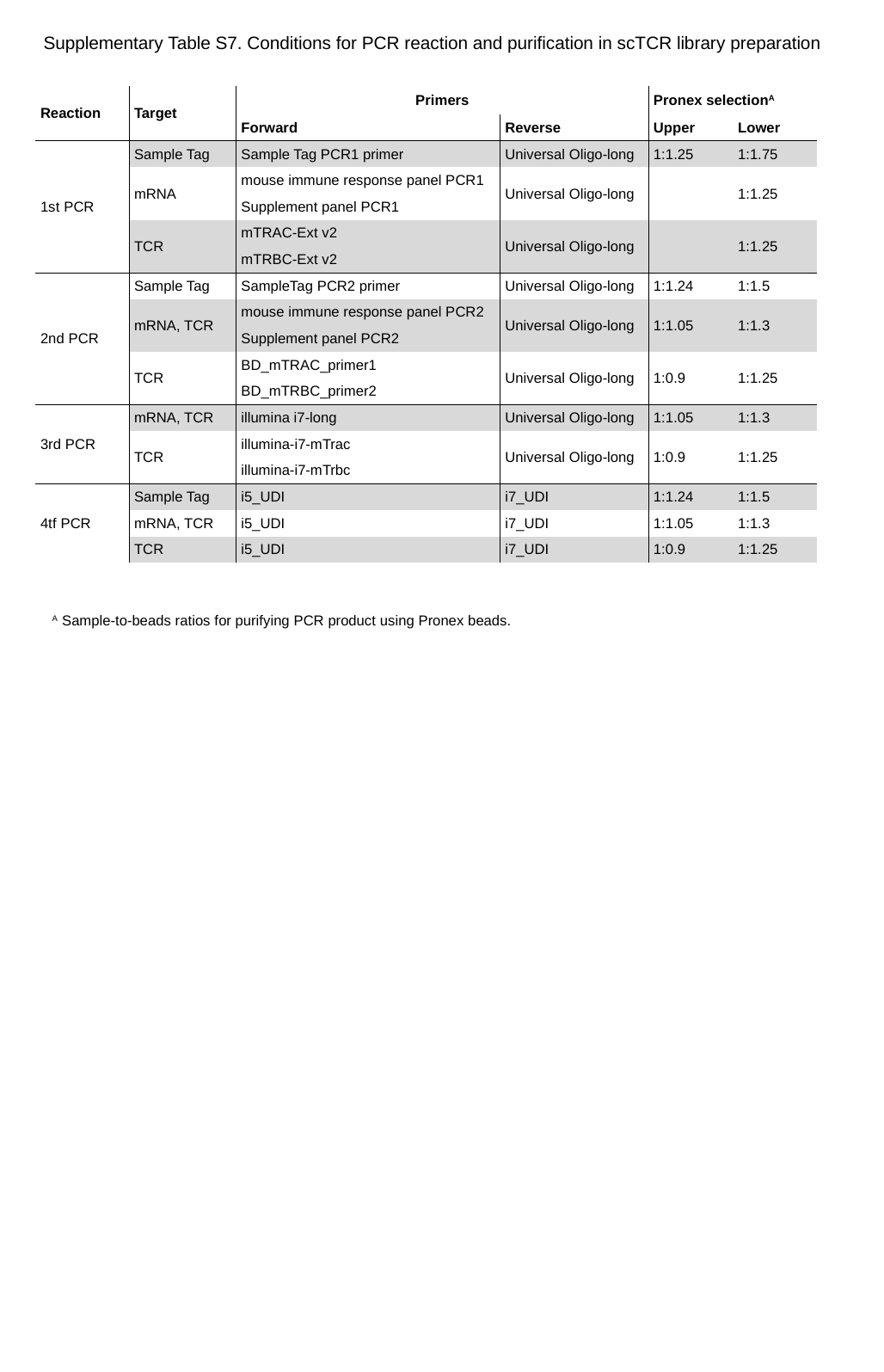

Supplementary Table S7. Conditions for PCR reaction and purification in scTCR library preparation
| Reaction | Target | Primers | | Pronex selection^A^ | |
| --- | --- | --- | --- | --- | --- |
| Reaction | Product | Forward | Reverse | Upper | Lower |
| 1st PCR | Sample Tag | Sample Tag PCR1 primer | Universal Oligo-long | 1:1.25 | 1:1.75 |
| | mRNA | mouse immune response panel PCR1 | Universal Oligo-long | | 1:1.25 |
| | | Supplement panel PCR1 | | | 1:1.25 |
| | TCR | mTRAC-Ext v2 | Universal Oligo-long | | 1:1.25 |
| | | mTRBC-Ext v2 | | | 1:1.25 |
| 2nd PCR | Sample Tag | SampleTag PCR2 primer | Universal Oligo-long | 1:1.24 | 1:1.5 |
| | mRNA, TCR | mouse immune response panel PCR2 | Universal Oligo-long | 1:1.05 | 1:1.3 |
| | | Supplement panel PCR2 | | | |
| | TCR | BD\_mTRAC\_primer1 | Universal Oligo-long | 1:0.9 | 1:1.25 |
| | | BD\_mTRBC\_primer2 | | | |
| 3rd PCR | mRNA, TCR | illumina i7-long | Universal Oligo-long | 1:1.05 | 1:1.3 |
| | TCR | illumina-i7-mTrac | Universal Oligo-long | 1:0.9 | 1:1.25 |
| | | illumina-i7-mTrbc | Universal Oligo-long | | |
| 4tf PCR | Sample Tag | i5\_UDI | i7\_UDI | 1:1.24 | 1:1.5 |
| | mRNA, TCR | i5\_UDI | i7\_UDI | 1:1.05 | 1:1.3 |
| | TCR | i5\_UDI | i7\_UDI | 1:0.9 | 1:1.25 |
^A^ Sample-to-beads ratios for purifying PCR product using Pronex beads.

### Slide 12
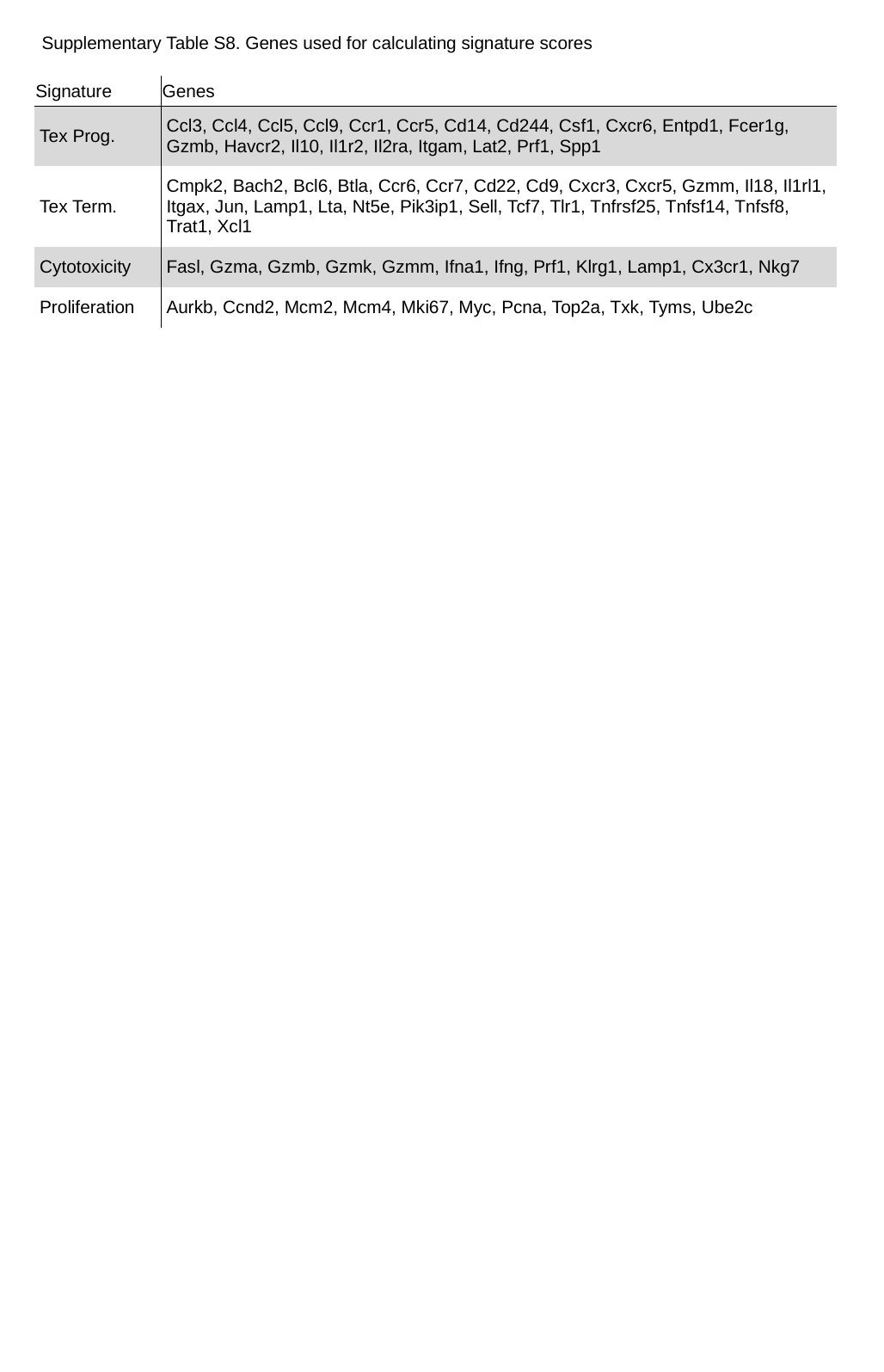

Supplementary Table S8. Genes used for calculating signature scores
| Signature | Genes |
| --- | --- |
| Tex Prog. | Ccl3, Ccl4, Ccl5, Ccl9, Ccr1, Ccr5, Cd14, Cd244, Csf1, Cxcr6, Entpd1, Fcer1g, Gzmb, Havcr2, Il10, Il1r2, Il2ra, Itgam, Lat2, Prf1, Spp1 |
| Tex Term. | Cmpk2, Bach2, Bcl6, Btla, Ccr6, Ccr7, Cd22, Cd9, Cxcr3, Cxcr5, Gzmm, Il18, Il1rl1, Itgax, Jun, Lamp1, Lta, Nt5e, Pik3ip1, Sell, Tcf7, Tlr1, Tnfrsf25, Tnfsf14, Tnfsf8, Trat1, Xcl1 |
| Cytotoxicity | Fasl, Gzma, Gzmb, Gzmk, Gzmm, Ifna1, Ifng, Prf1, Klrg1, Lamp1, Cx3cr1, Nkg7 |
| Proliferation | Aurkb, Ccnd2, Mcm2, Mcm4, Mki67, Myc, Pcna, Top2a, Txk, Tyms, Ube2c |

### Slide 13
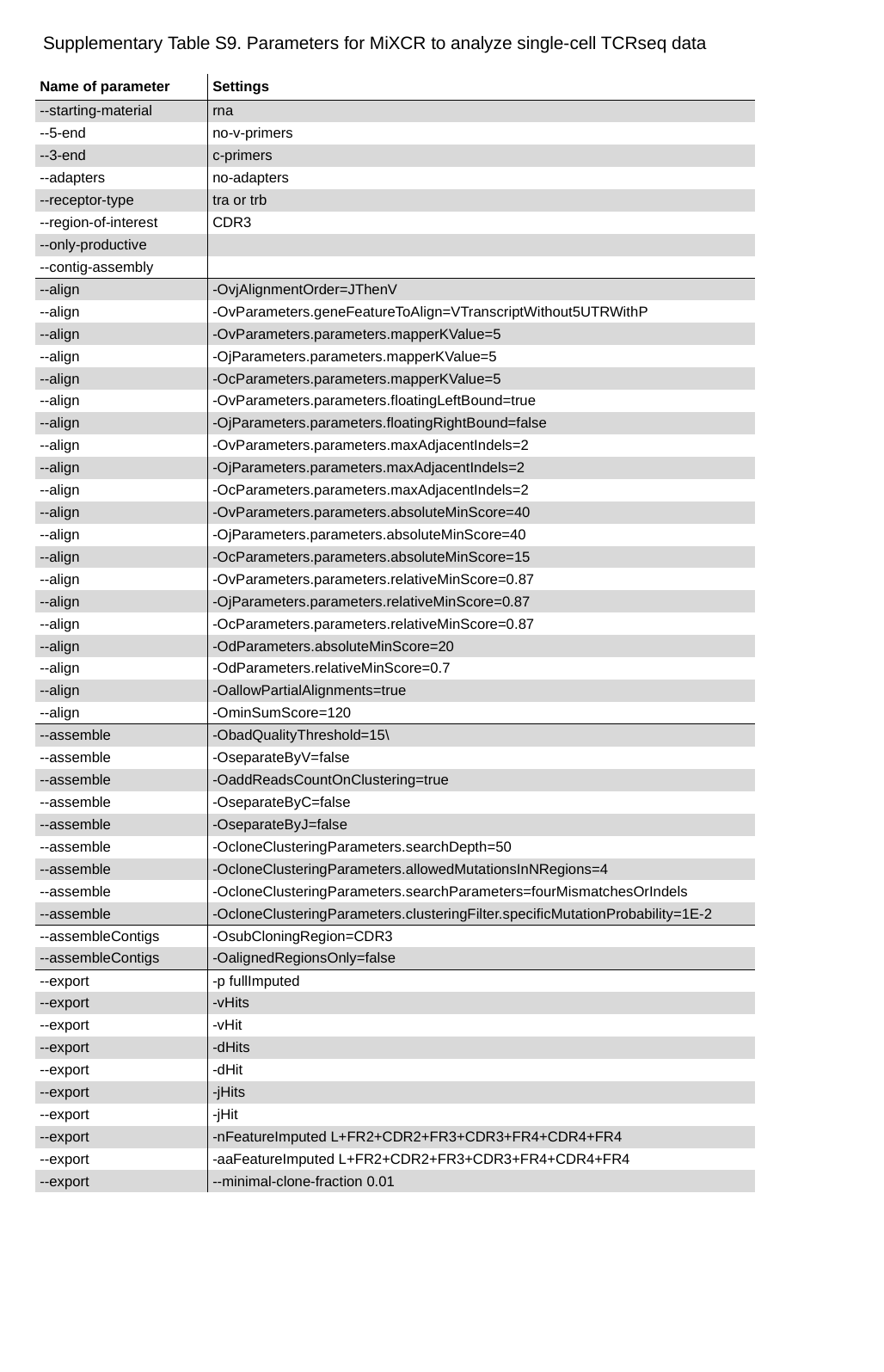

Supplementary Table S9. Parameters for MiXCR to analyze single-cell TCRseq data
| Name of parameter | Settings |
| --- | --- |
| --starting-material | rna |
| --5-end | no-v-primers |
| --3-end | c-primers |
| --adapters | no-adapters |
| --receptor-type | tra or trb |
| --region-of-interest | CDR3 |
| --only-productive | |
| --contig-assembly | |
| --align | -OvjAlignmentOrder=JThenV |
| --align | -OvParameters.geneFeatureToAlign=VTranscriptWithout5UTRWithP |
| --align | -OvParameters.parameters.mapperKValue=5 |
| --align | -OjParameters.parameters.mapperKValue=5 |
| --align | -OcParameters.parameters.mapperKValue=5 |
| --align | -OvParameters.parameters.floatingLeftBound=true |
| --align | -OjParameters.parameters.floatingRightBound=false |
| --align | -OvParameters.parameters.maxAdjacentIndels=2 |
| --align | -OjParameters.parameters.maxAdjacentIndels=2 |
| --align | -OcParameters.parameters.maxAdjacentIndels=2 |
| --align | -OvParameters.parameters.absoluteMinScore=40 |
| --align | -OjParameters.parameters.absoluteMinScore=40 |
| --align | -OcParameters.parameters.absoluteMinScore=15 |
| --align | -OvParameters.parameters.relativeMinScore=0.87 |
| --align | -OjParameters.parameters.relativeMinScore=0.87 |
| --align | -OcParameters.parameters.relativeMinScore=0.87 |
| --align | -OdParameters.absoluteMinScore=20 |
| --align | -OdParameters.relativeMinScore=0.7 |
| --align | -OallowPartialAlignments=true |
| --align | -OminSumScore=120 |
| --assemble | -ObadQualityThreshold=15\ |
| --assemble | -OseparateByV=false |
| --assemble | -OaddReadsCountOnClustering=true |
| --assemble | -OseparateByC=false |
| --assemble | -OseparateByJ=false |
| --assemble | -OcloneClusteringParameters.searchDepth=50 |
| --assemble | -OcloneClusteringParameters.allowedMutationsInNRegions=4 |
| --assemble | -OcloneClusteringParameters.searchParameters=fourMismatchesOrIndels |
| --assemble | -OcloneClusteringParameters.clusteringFilter.specificMutationProbability=1E-2 |
| --assembleContigs | -OsubCloningRegion=CDR3 |
| --assembleContigs | -OalignedRegionsOnly=false |
| --export | -p fullImputed |
| --export | -vHits |
| --export | -vHit |
| --export | -dHits |
| --export | -dHit |
| --export | -jHits |
| --export | -jHit |
| --export | -nFeatureImputed L+FR2+CDR2+FR3+CDR3+FR4+CDR4+FR4 |
| --export | -aaFeatureImputed L+FR2+CDR2+FR3+CDR3+FR4+CDR4+FR4 |
| --export | --minimal-clone-fraction 0.01 |
